# Long-term adaptation to hypoxia provides insight into mechanisms facilitating the switch of *Pseudomonas aeruginosa* to chronic lung infections

**DOI:** 10.1101/2024.12.26.630387

**Authors:** Joanna Drabinska, Lucia O’Connor, Niamh Duggan, Ciarán J. Carey, Siobhán McClean

## Abstract

Opportunistic bacterial infections are an increasing threat, especially for immunocompromised individuals such as people with cystic fibrosis (CF), driving morbidity and mortality. *Pseudomonas aeruginosa* is a key pathogen associated with chronic lung infections, that has been extensively studied in this context, but the mechanism(s) driving its adaptation towards chronic colonisation in the lung are not fully understood. This work focuses on the adaptations of *P. aeruginosa* to long-term hypoxia, one of the important environmental pressures present in the CF lung, to investigate whether it drives the development of persistence in CF patients. We used an experimental evolution approach to investigate how an early CF strain adapted to 6% oxygen over 28 days. We focussed on the impact of long-term hypoxia on the proteome and investigated the emergence of stable changes in phenotype. Changes in the abundance of >140 proteins were observed compared to the ancestral strain, including proteins involved in antibiotic resistance, stress response, iron homeostasis, biofilm formation and those previously associated with chronic infection. Significant changes in the abundance of proteins regulating cellular c-di-GMP levels were also observed. We show that two distinct *P. aeruginosa* small colony variants (SCVs) emerged, one exclusively in hypoxia exposed cultures. Hypoxia-adapted cultures developed resistance to 8 out of 13 antibiotics tested; increased biofilm and exopolysaccharide production; and decreased pyocyanin production, consistent with the changes in the proteome. All hypoxia-adapted cultures showed decreased siderophore production. Overall, we demonstrate that long-term hypoxia exposure contributes to multiple changes in phenotype and proteome that are frequently observed in *P. aeruginosa* CF lung chronic infection isolates. This suggests that hypoxia is driving these adaptations, at least in part, and opens a new path to treatment.

**Author Summary:** Opportunistic antibiotic-resistant bacterial infections are an increasing threat, particularly for immunocompromised individuals such as people with cystic fibrosis (CF). These infections increase disease severity, the number of deaths and healthcare costs. *Pseudomonas aeruginosa* is a major pathogen in this context, causing persistent infections, yet the mechanisms driving its adaptation towards chronic colonisation in the CF lung are not understood.

We investigated the adaptations that *P. aeruginosa* exposed to low oxygen conditions (an important environmental pressure in the CF lung) for 28 days, to examine whether it drives adaptations that lead to chronic infection. We found that prolonged exposure to hypoxia induced stable changes in *P. aeruginosa*, which are distinct from those reported after short term exposure. We observed multiple adaptations in hypoxia adapted cultures, many of which were associated with important signalling networks linked to lifestyle shifts in bacteria and several were also observed in isolates that had adapted to chronic infection. Similar adaptive responses in other opportunistic environmental pathogens suggest that this process of adaptation could be targeted to create universal therapeutic approaches preventing antibiotic overuse and improving patient outcomes.

## Introduction

Antibiotic-resistant bacterial infections are a growing threat to society. They were associated with nearly five million deaths in 2019 (1), and are expected to cause more than 10 million deaths annually by 2050 (2). Opportunistic pathogens such as *Pseudomonas aeruginosa* play a crucial role in escalating this crisis. While these bacteria typically do not cause infections in healthy people, immunocompromised individuals and people with underlying conditions such as cystic fibrosis (CF) and chronic obstructive pulmonary disease (COPD) or those on ventilator support are at great risk of developing chronic *P. aeruginosa* lung infections (3,4). Moreover, opportunistic pathogens are responsible for the majority of healthcare-associated infections, contributing to increasing morbidity, mortality and costs (5).

*P. aeruginosa* is a Gram-negative opportunistic human pathogen, which is a leading cause of nosocomial infections, constituting 11-26% of ventilator-associated pneumonia globally (6). Overall, it causes more than 7% of all health-associated infections and is a major contributor to morbidity and mortality among people with CF (7–9). It has many intrinsic and acquired mechanisms of antibiotic resistance contributing to its spread and persistence (10), and the prolonged antibiotic treatments necessary to manage *P. aeruginosa* chronic infections contribute to the spread of antimicrobial resistance (11,12). Furthermore, the World Health Organisation recognised *P. aeruginosa* as a high priority pathogen in 2024 while the European Centre for Disease Prevention and Control (ECDC) considers it a serious threat (13–16).

A key feature of *P. aeruginosa* is its exquisite metabolic flexibility and adaptability to harsh conditions, allowing it to transition from the colonisation of the rhizosphere or hydrosphere to colonising niches within human or animal hosts, and establishing persistent infections (4,17). *P. aeruginosa* chronic infection isolates show multiple changes that attenuate virulence, allowing it to avoid immune recognition, thereby enhancing survival, and consequently, chronic infections are extremely difficult to eradicate (18). Its adaptability is driven by a complex regulatory network involving transcriptional regulators, DNA modifications, non-coding RNAs, quorum sensing, and a range of virulence factors (18–23). These adaptations and associated lifestyle changes are also driven by the alterations in c-di-GMP second messenger levels (24–27). Despite this extensive research, the mechanism(s) enabling *P. aeruginosa* to transition from acute to chronic infection remain unclear.

*P. aeruginosa*, a facultative anaerobe, thrives under low oxygen and anaerobic conditions using alternative terminal electron acceptors and fermentation substrates. It preferentially colonizes hypoxic niches within mucus layers of the lung over more oxygenated surfaces (28). *P. aeruginosa* responds to changes in oxygen availability via the transcriptional regulator Anr, which regulates many genes shown to be altered in CF chronic infection (29,30). Studies on another CF-associated pathogen, *Burkholderia cenocepacia* have shown that 40% of proteins encoded within a low oxygen activated (Lxa) locus showed increased abundance in chronic infection isolates from CF patients (31). Subsequent studies showed that deletion of single genes from the Lxa or the entire locus resulted in changes in phenotypes crucial for the virulence of this bacterium including host cell attachment, intracellular survival and stress responses (31–33). Additionally, the *sicX* sRNA has been proposed to be a biomarker of *P. aeruginosa* chronic infection and is also oxygen responsive (34). Given that hypoxia is an important environmental pressure in the CF lung, we wanted to evaluate the impact of long-term hypoxia exposure on *P. aeruginosa*.

We developed an experimental evolution approach and demonstrate that the impact of long-term exposure to low oxygen on the proteome and phenotype of *P. aeruginosa* is consistent with many adaptations that are associated with chronic lung infection. We specifically evaluated on the proteome and phenotype to focus on the effective changes in the strains following long-term hypoxia, which is not widely studied. Our findings provide new insights into the mechanisms that underlie the persistence and pathogenesis of *P. aeruginosa* in chronic infection and possibly other opportunistic pathogens, within the host.

## Results

### Design and Optimisation of Experimental Adaptation

To investigate the impact of long-term exposure to low oxygen conditions on the adaptation of *P. aeruginosa*, we designed an experimental evolution/adaptation approach (Fig 1A), exposing the *P. aeruginosa* early infection strain AMT 0023-30 to either 6% oxygen or to normoxia (21%) for 28 days. AMT 0023-30 is a well-characterised and sequenced CF paediatric early infection strain that was selected from the international *P. aeruginosa* panel (35–38). This strain was isolated from a 6 month old paediatric patient with no former history of *P. aeruginosa* infection, thus the likelihood of preadaptation to the lung or hypoxic niches is very low. Furthermore, a sequential, 96-month, chronic infection strain is available (AMT 0023-34) which has also been sequenced and well characterised (35–38). This strain is the ancestor of the adapted cultures further described in this manuscript (Table 1). After considering the ranges of oxygen that *P. aeruginosa* is exposed to in soil and water, together with the range of hypoxic conditions reported in the CF lung and available literature on physiological tissue hypoxia (28,39,40), we chose 6% oxygen as a low oxygen concentration. Soil aeration ranges from 10 to 20% but decreases with depth and microoxic and anoxic niches are available (39). Similarly a range of oxic, hypoxic and anoxic conditions have been reported for CF patient lungs (28, 40) ranging from180 mmHg in the lumen to 2.5 mmHg within the mucus. Thus 6% O_2_ is in the higher hypoxia range and it allowed us to observe the response to hypoxia, while not excessively stressing the cells, so that we can distinguish this response from changes associated with anoxia.

**Figure 1.**
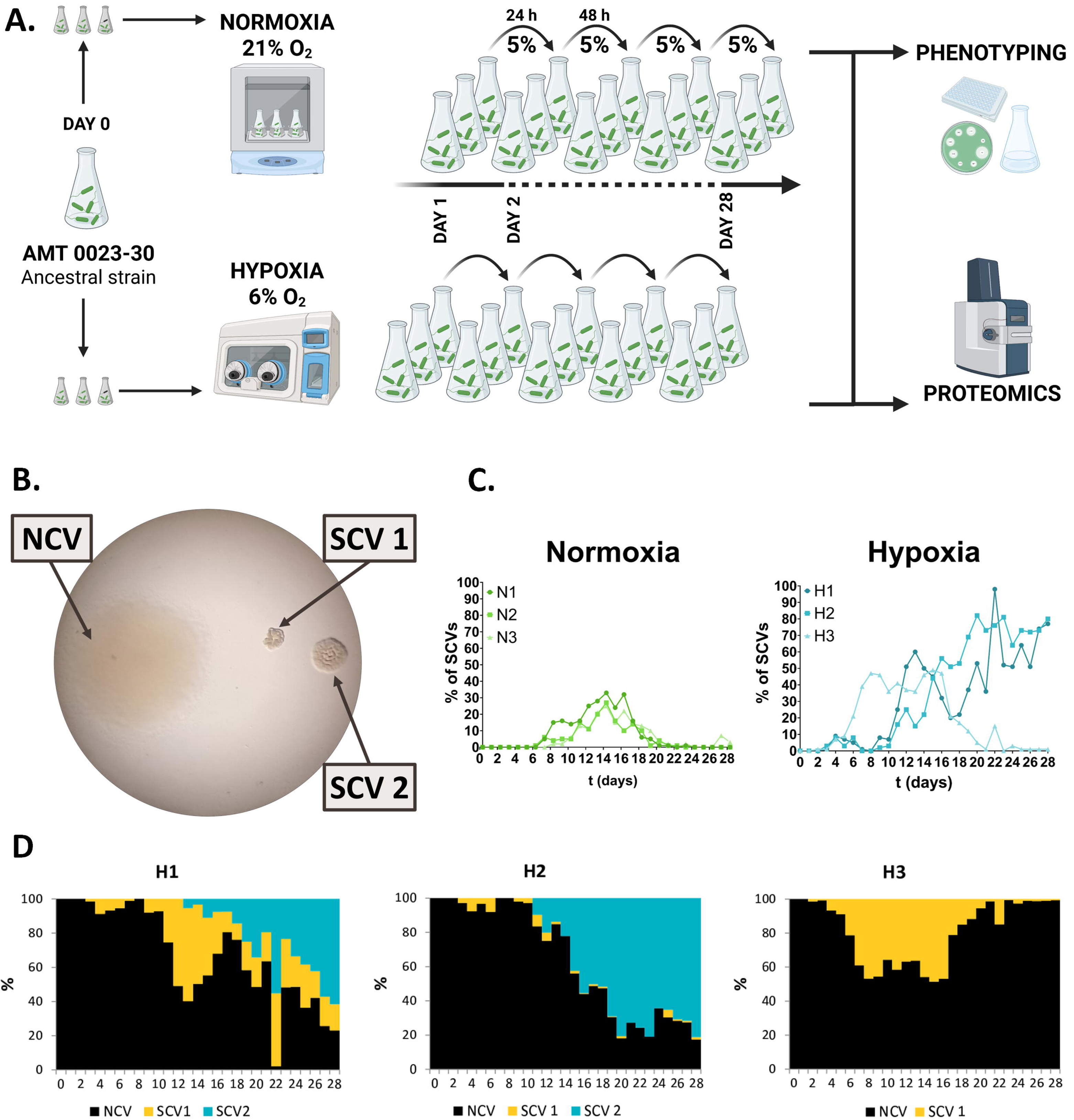
Long-term adaptation of Pseudomonas aeruginosa to hypoxic conditions. Experimental evolution study design (A.) and influence of hypoxia on small colony variants emergence (B.-D.). A. Experimental evolution study design. The overnight culture of the ancestral strain was inoculated into three flasks with hypoxia or normoxia equilibrated medium and incubated with agitation in 6% O_2_ or 21% O_2_. Every 24 h 5% of the culture was transferred into a fresh portion of equilibrated media. The transfers of the hypoxia-adapting cultures were performed under hypoxic conditions to avoid oxygen level fluctuation. The samples from the cultures adapted for 28 days were taken for further proteomic and phenotypic analysis. The details of the procedure are described in the method section. B. Hypoxia-adapted colony morphotypes observed throughout the long-term adaptation. C. Percentage of SCVs in individual normoxia- and hypoxia adapting cultures over 28 days. D. Ratios of SCV 1 and SCV 2 in the hypoxia-adapting populations over 28 days. The percentage of the small colony variants in the whole populations was carried out for each flask individually each day.

**Table 1.** List of strains and populations used in this study.

| Strain/Population Name |  | Description | Source |
| --- | --- | --- | --- |
| E |  | Paediatric (6-month old) early infection cystic fibrosis strain AMT 0023-30. Ancestral strain. | (38) |
| L |  | Paediatric late infection cystic fibrosis strain AMT 0023-34, sequential (96 months) isolate of AMT 0023-30. | (38) |
| NACs | N1 | 28 day normoxia-adapted population, replicate 1. Evolved from AMT 0023-30 (E). | This study |
|  | N2 | 28 day normoxia-adapted population, replicate 2. Evolved from AMT 0023-30 (E). | This study |
|  | N3 | 28 day normoxia-adapted population, replicate 3. Evolved from AMT 0023-30 (E). | This study |
| HACs | H1 | 28 day hypoxia-adapted population, replicate 1. Evolved from AMT 0023-30 (E). | This study |
|  | H2 | 28 day hypoxia-adapted population, replicate 2. Evolved from AMT 0023-30 (E). | This study |
|  | H3 | 28 day hypoxia-adapted population, replicate 3. Evolved from AMT 0023-30 (E). | This study |

To maintain stable growth over the period of the evolution experiment, daily culture transfer volumes were optimised by evaluating a range of volumes (1-10%) of overnight *P. aeruginosa* cultures transferred into fresh media and analysing their growth patterns (S1 Figure). A 5% volume transfer was chosen as optimal, as it allowed the transfer of the maximum number of bacterial cells without disrupting the growth pattern and resulted in a starting OD_600_ of ∼0.1. Examination of growth patterns of the early infection strain revealed small differences in the generation times observed during hypoxic and normoxic conditions, with the strain growing 1.8-fold slower under hypoxia (g = 107 minutes) compared to normoxia (g = 60 minutes), and showing logarithmic phases of 3 h in normoxia and 6 h in hypoxia (S2 Figure). Consequently, we estimate that during the 28 day adaptation period, all the cultures underwent 84-94 generations.

To facilitate further investigation, including phenotyping and proteome analysis, three independent flasks were inoculated and incubated under either 6% oxygen or normoxia (21% O_2_), and samples were collected every day over 28 days, at the time of culture transfer. Three replicate cultures were chosen for each condition to facilitate deeper analysis over a broad range of phenotypes. To maintain stable hypoxic conditions and avoid exposure to fluctuating oxygen levels, the transfers of hypoxia-adapting cultures were conducted within the hypoxic chamber. The three 28 day independently adapted populations evolved from the ancestral AMT 0023-30 strain are referred to as N1, N2, and N3 for replicate normoxia-adapted control populations (NACs) and H1, H2, and H3 for the three replicate hypoxia-adapted cultures (HACs) (Table 1). All characterisation of the individual adapted populations (both HACs and NACs) was performed under normoxic conditions, to ensure the observed changes were stable in the absence of continued hypoxia and are not a transient, plastic response.

### Changes in colony morphology and pyocyanin production during adaptation

Changes in colony morphology often serve as key indicators of adaptation to the environment. Plating of daily samples from the adapting cultures showed the emergence of small colony variants (SCVs) in both NACs and HACs; however, the SCVs appeared earlier in HACs over the timeframe (Fig 1B and 1C). SCVs first appeared on day 2 in the H3 population, and by day 3 in the H1 and H2 populations. The SCV abundance peaked at 98% on day 22, with 77% remaining on day 28 in H1. A similar profile was shown in H2, with SCVs comprising 82% of the population by day 20. In contrast in population H3, the SCV abundance was generally lower and more variable, reaching only 50% of the population by day 15, and falling to 1% on day 21 (Fig 1C). The SCVs appeared much later in normoxic cultures, on days 6, 7 and 8 in flasks N1, N2 and N3 respectively, and only reached a maximum of 25-33% of the population by day 14 (Fig 1C).

Two distinct SCV morphologies were observed (Fig 1B and 1D). The first type, termed SCV 1, was observed both under normoxia and hypoxia, and resembled the morphology most often described as a rugose small colony variant characterised by small size, irregular edges and a heavily wrinkled, dry surface (41). The second type, SCV 2, was bigger than SCV 1 with a similar rough, dry surface; however it was easily distinguished by a more regular edge and shape and importantly, it was exclusive to HACs. The SCVs were stable after replating in normoxic conditions, although SCV 2 morphology differed, with a smoother surface and edges (S3 Figure).

Throughout the 28 days of adaptation, we also observed dynamic changes in culture colour (Fig 2A), suggesting changes in pigments, such as pyocyanin. Despite almost no visible change in colour being observed on day 28 (Fig 2A), quantification of pyocyanin production revealed that populations H1, H2 and one NAC (N1) produced comparable pyocyanin levels at day 28 to the late infection strain (AMT 0023-34) (Fig 2B and Fig 2C). Although pyocyanin was higher in the NACs overall compared with HACs, the differences between biological replicates suggests that pyocyanin production is independent of environmental oxygen levels or is not stable in normoxic conditions.

**Figure 2.**
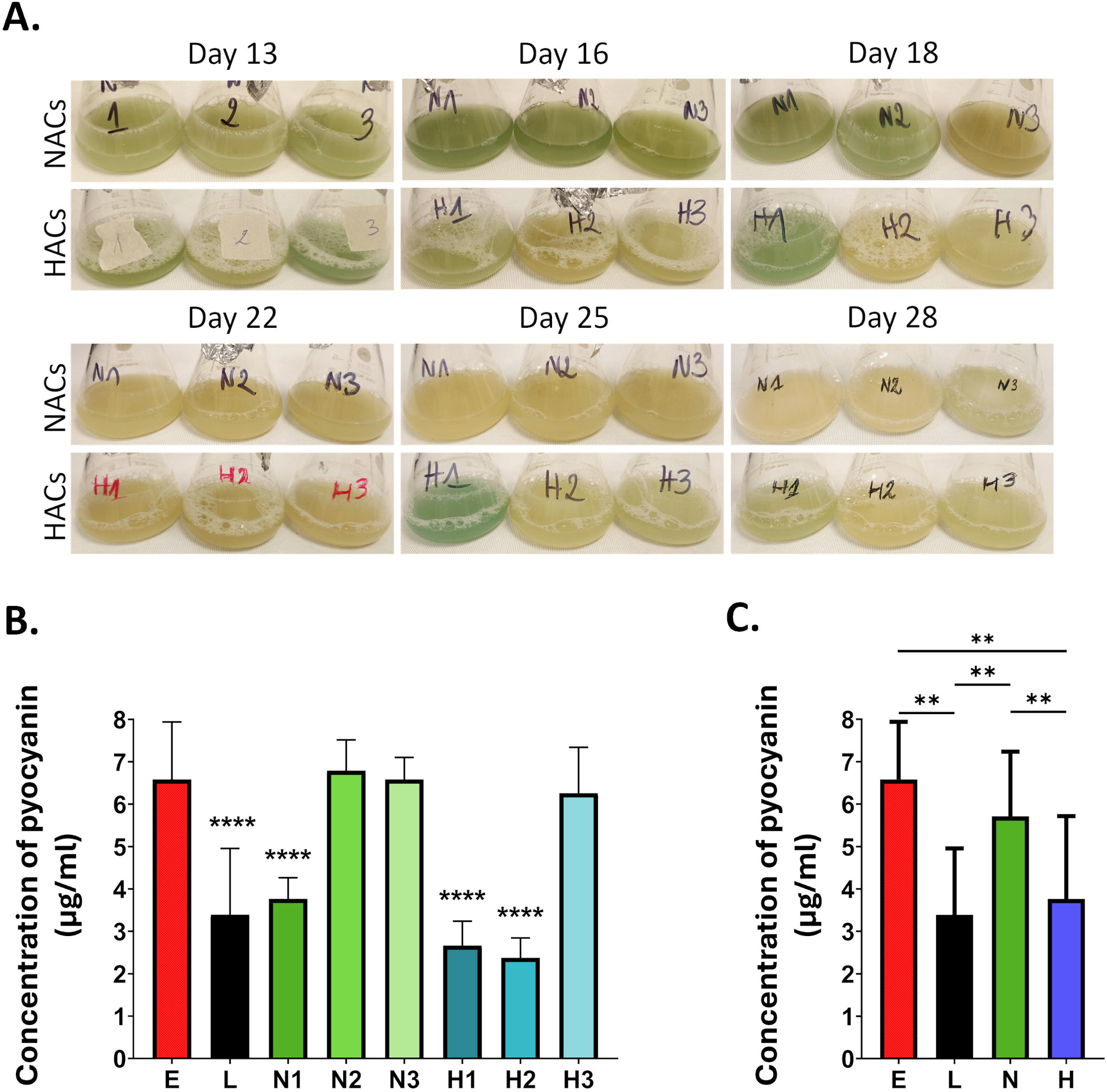
Influence of hypoxia on pyocyanin production. A. Representative photographs of flasks during the 28 day experimental evolution study showing changes in pyocyanin production. B. and C. Quantification of pyocyanin production by individual populations (B.) and mean values for normoxia- and hypoxia-adapted cultures (C.) at 28 days, relative to early (E) and late (L) infection strains. The data represent the concentration of pyocyanin in 1ml of bacterial culture, calculated as described in the methods. The data represent two independent experiments performed in normoxic conditions, with at least three technical replicates for each strain. Statistically significant differences were determined by ordinary one-way ANOVA test compared to the early infection strain with an uncorrected Fishers LSD test (B.) or Kruskal-Wallis one-way ANOVA test comparing all groups with an uncorrected Dunn’s test (C.) are shown (**p< 0.01; ***p<0.001; ****p< 0.0001). E: early infection clinical strain; L: late infection clinical strain; N1, N2, N3: 28-day normoxia-adapted populations; H1, H2, H3: 28-day hypoxia adapted populations.

### Influence of hypoxia on the whole proteome

The proteomes of 28 day adapted cultures were analysed to elucidate the overall influence of long-term hypoxia on *P. aeruginosa*. The individual flasks were treated as biological replicates, thus showing the global changes across three independent populations. Principal component analysis (PCA) showed a clear adaptation to the conditions tested, PC1 and PC2 accounted for 84.0% and 7.3% of the total variance respectively (Fig 3A). A total of 257 proteins showed significant differences (adjusted p-value ≤0.10) in abundance between HACs and the ancestor strain, NACs and the ancestor strain and/or between HACs and NACs (Table 2, S4 Table). The majority of proteins that were altered in abundance between HACs, and the ancestral strain were decreased following exposure to hypoxia (n=106; 72.55%), while only 36 proteins (27.45%) showed increased abundance (Table 2, volcano plot, Fig 3C). An additional 13 proteins were identified as significantly altered in abundance after comparing HACs to NACs. Eighty proteins were changed both in HACs and NACs relative to the ancestral strain and, suggesting they may be affected by long-term culture generally and not hypoxia. A total of 89 proteins were further considered as significantly altered during hypoxia adaptation (Fig 3B).

**Figure 3.**
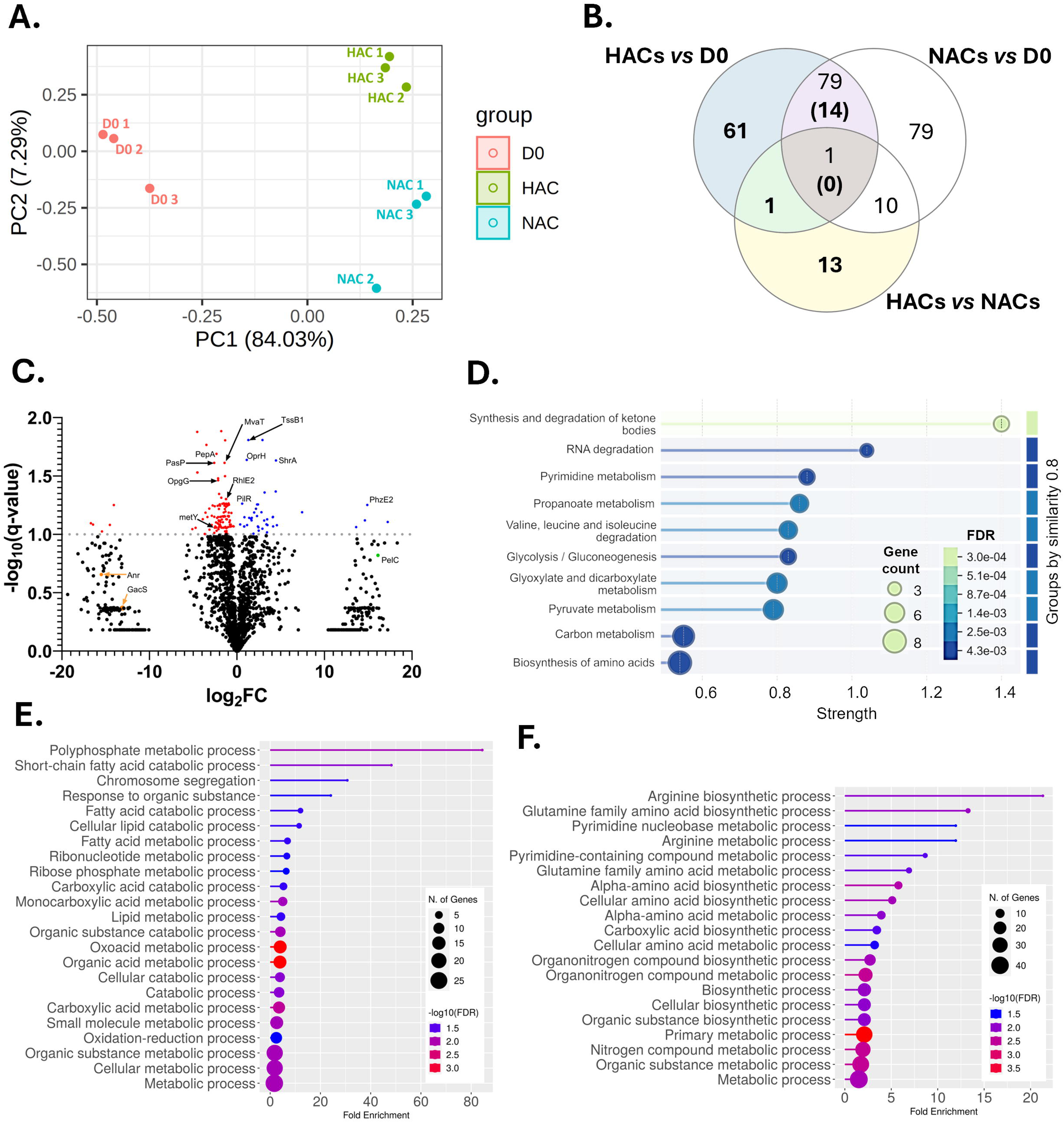
Whole proteome comparison of long-term hypoxia-adapted and normoxia-adapted P. aeruginosa. A. Principal component analysis plot of quantified proteins from label-free LC-MS/MS analysis of whole cell extracts from P. aeruginosa adapted to hypoxic conditions (HACs) or normoxic conditions (NACs) for 28 days and the ancestral strain (D0). The PCA plot was performed using LFQ intensities of 1785 proteins across three biological replicates per condition. Clustering by condition indicates specific global proteomic shifts associated with long-term hypoxia adaptation. The PCA plot was generated using the Galaxy server v 3.4.0 (42) with ggplot2 using default settings with Euclidean distance clustering. B. A Venn diagram of the number of proteins significantly changed in abundance between HACs and the ancestral strain; NACs and the ancestral strain; and HACs vs NACs. The analysis was performed using one-way ANOVA with Benjamini-Hochberg correction and proteins with an adjusted p-value <0.1 were considered significantly altered in abundance. The 89 proteins in the coloured sections were considered for further analysis. The white sections were not considered due to lack of relevance to hypoxia adaptation. Only the numbers in parenthesis in the cross section of NACs and HACs compared to the ancestral strains were considered due to distinct change between HACs and NACs (1.5 fold-change). C. Volcano plot of differentially expressed proteins between HACs vs the ancestral strain (D0) representing log_10_ of adjusted p-values and log_2_ transformed fold-change values. Proteins significantly decreased in abundance are highlighted in red and proteins significantly increased in abundance are highlighted blue. Examples of proteins of interest are labelled. A value of 1 was imputed for all the zero LFQ values, thus they are represented on the plot, despite the lack of detection. D. STRING-based KEGG pathway enrichment of 89 proteins significantly altered in abundance in HACs vs the ancestral strain and HACs vs NACs. Enrichment analysis was conducted using STRING functional enrichment (FDR <0.05) (v12.0) (43). The dot plot shows the top 10 enriched pathways, where dot size corresponds to the number of hits per pathway and colour represents statistical significance (FDR-corrected p-value). The pathways are grouped by similarity (0.8). E. and F. Dot plots representing functional GoTerm Biological Process enrichment of proteins with increased (E) and decreased abundance (F). Size corresponds to the number of hits per pathway and colour represents statistical significance. The enrichment analysis and dot plot were prepared using ShinyGo v0.82 (44).

**Table 2.**
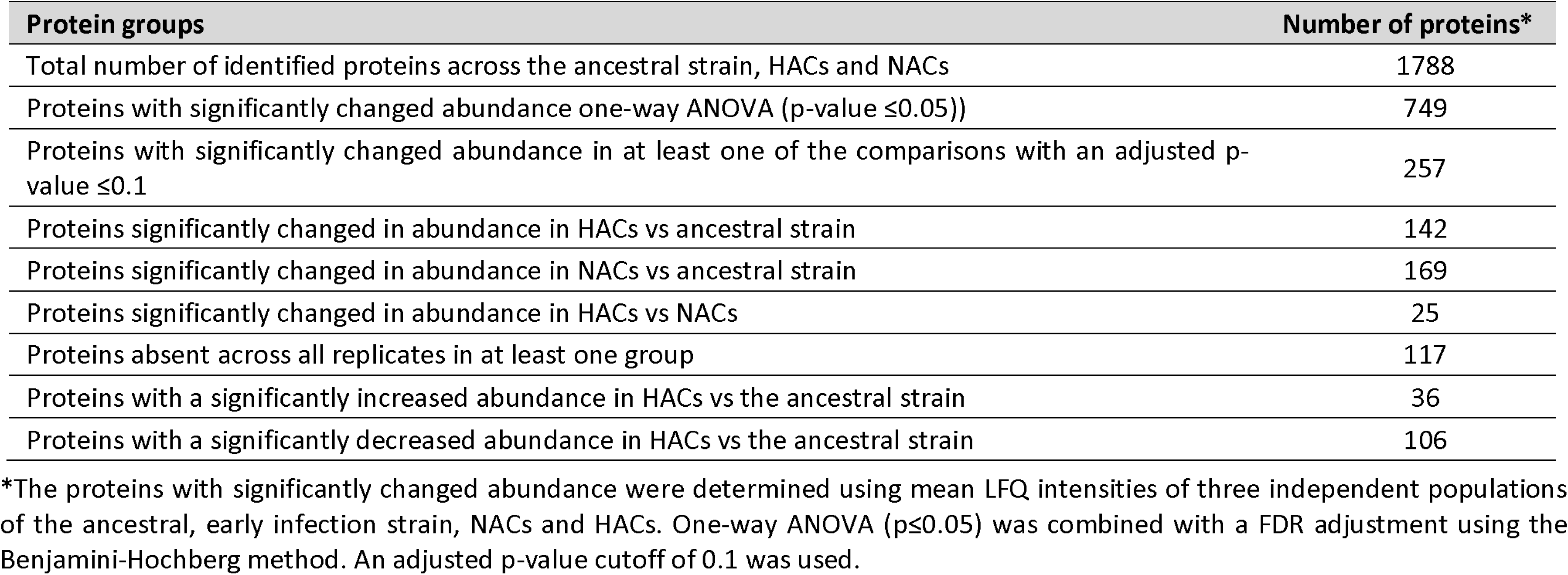
Proteome comparison of 28 day the P. aeruginosa HACs and NACs vs the early infection ancestral strain and HACs vs NACs.

### Proteins unique to hypoxia or normoxia

Interestingly, 117 (15.62%) of the 749 proteins that showed significant changes in abundance (one-way ANOVA) were undetectable in at least one of the groups: all three HACs, all three NACs or all three ancestral strain replicates. Their consistent presence or absence suggests that their expression was either switched on or off, and consequently, they are likely to play critical roles under the specific conditions. Among these, only three proteins were unique to HACs and not detected in any NAC or in the ancestral strain: phenazine biosynthesis protein, PhzE; Pel polysaccharide secretion protein, PelC; and glutamine amidotransferase, HisH2 (Table 3). Additionally, anthranilate synthase component I PhnA and iron(II)/α-ketoglutarate-dependent oxygenase AmbC, involved in toxin production were undetected in the ancestral strain, and expressed in all three HACs, but only one of the NACs.

**Table 3.**
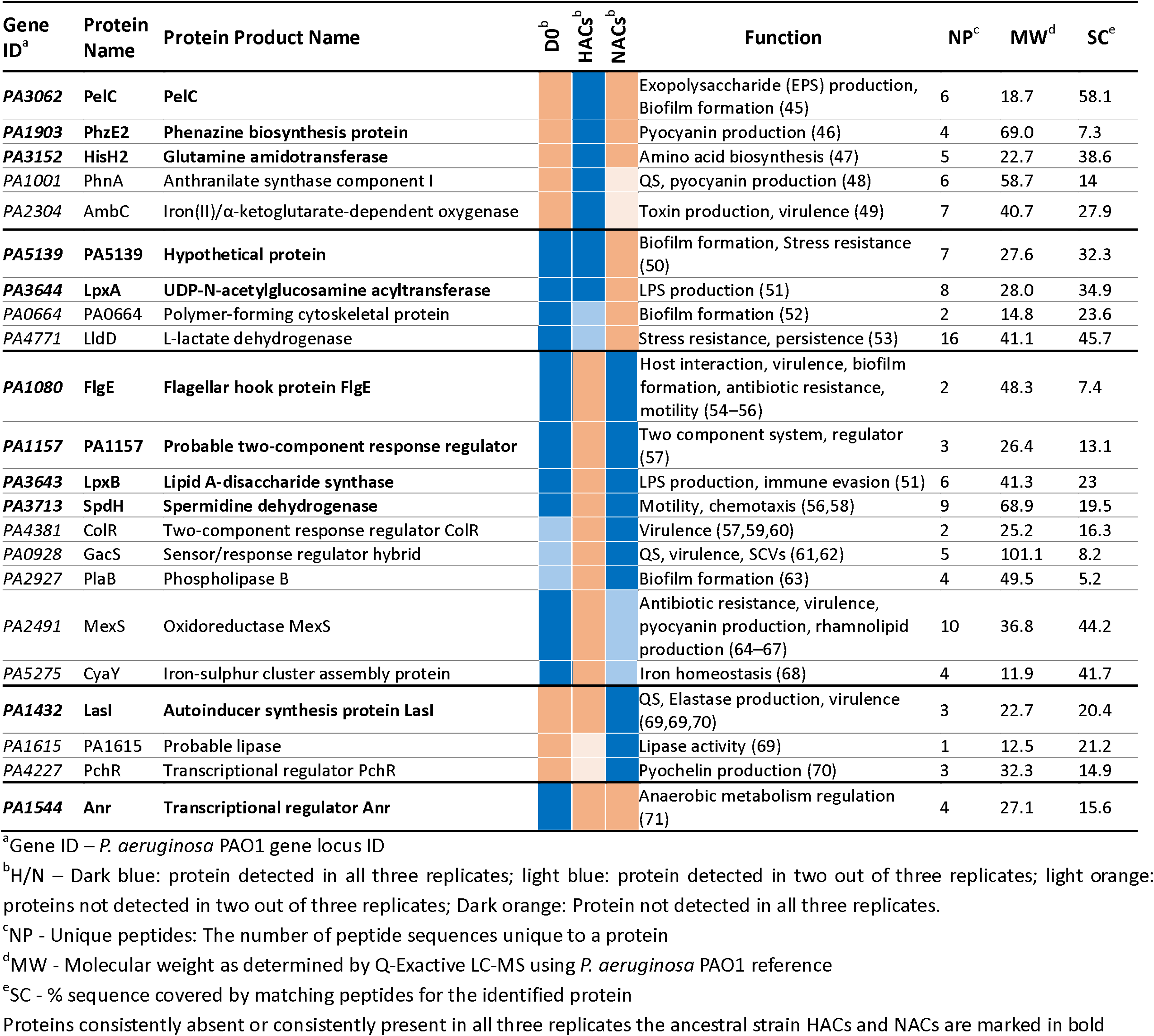
Examples of proteins not detected in at least one of the groups (Day 0, HACs, NACs).

Many proteins that were undetectable in HACs but expressed in the ancestral strain and all NACs and thus potentially redundant in low oxygen conditions, are involved in virulence, including flagellar hook protein FlgE, involved in motility, virulence, host interactions, biofilm formation and antibiotic resistance; lipid A-disaccharide synthase LpxB, involved in immune evasion; spermidine dehydrogenase SpdH, involved in chemotaxis and motility. In addition, GacS and ColR, components of two-component response regulator systems involved in virulence regulation, and phospholipase B were also undetectable in all HACs, but were present in two of the replicates of the ancestral strain, and all three NACs. Moreover, oxidoreductase MexS associated with antibiotic resistance and virulence factor production, chromosome partitioning protein MksE, and iron-sulphur cluster assembly protein CyaY were detected in all replicates of the ancestral strain and two of the NACs, while undetectable in all three HACs (Table 3). This attenuation of virulence proteins mirrors the loss of virulence during chronic infection.

Proteins which were detected both in the ancestral strain and HACs but not detected in NACs may also be important in the process of adaptation to hypoxia, as their abundance is maintained. The transcription termination factor Rho associated with biofilm formation and stress resistance, and UDP-N-acetylglucosamine acyltransferase LpxA belong to this group. Additionally, polymer-forming cytoskeletal protein PA0664 involved in biofilm formation, and L-lactate dehydrogenase LldD were detected in all replicates of the ancestral strain and two of the HACs, but none of the NACs, suggesting their maintenance is required for fitness under low oxygen. Lastly, the proteins which were undetected in the ancestral strain replicates and all HACs include the acyl-homoserine lactone synthase LasI which produces a quorum-sensing signal and positively regulates numerous *P. aeruginosa* virulence-associated traits including biofilm formation, oxidative stress resistance and elastase production. Additionally, lipase PA1615 and transcriptional regulator PchR, controlling pyochelin production were not detected in all replicates of the ancestral strain and in two of the HACs while expressed in all three NACs. Interestingly, the Anr transcriptional regulator was undetectable both in normoxic and hypoxic conditions.

### Functional enrichment

Functional enrichment analysis of the 89 proteins identified in the previous step (Fig 3B) was performed. GO biological process identified 53 enriched categories (S5 Table), including various processes associated with carbon, nitrogen and amino acid metabolism; fatty acid and lipid metabolism; nucleotide and ribonucleotide metabolism. A separate analysis of proteins with increased or decreased abundance in HACs provided further insights into these processes (S5 Table). Overall, 25 GO biological processes were enriched among proteins increased abundance in hypoxia (Fig 3E), while 27 processes were enriched in the decreased abundance group (Fig 3F). Hypoxia led to upregulation of processes related to ribonucleotide metabolism, but a downregulation of pyrimidine-containing compound metabolism. Additionally, processes associated with lipid and fatty acid metabolism, oxidation-reduction and chromosome segregation were upregulated in HACs. Multiple processes associated with amino acid metabolism were also downregulated including glutamine and arginine. Additionally, nitrogen metabolism associated processes were downregulated as well. KEGG pathway enrichment analysis further highlighted alterations of pyrimidine metabolism, changes of RNA degradation pathways, and various carbon, amino acid, and energy metabolism associated pathways (Fig 3D).

### Antibiotic resistance of adapted cultures

A change in the abundance of many proteins associated with the resistance to multiple classes of antibiotics was observed in HACs (Table 4). Six proteins were increased in abundance including UDP-N-acetylmuramate-alanine ligase MurC, outer membrane protein H1 precursor OprH, and the Topoisomerase IV subunit A ParC, while five were decreased in abundance.

**Table 4.** Examples of proteins altered in abundance in P. aeruginosa in three comparisons.

| Gene ID<sup>a</sup> | Protein Name | Protein Product Name | Fold-change<sup>b</sup> |  |  | Function<sup>c</sup> | NP<sup>d</sup> | MW<sup>e</sup> | SC<sup>f</sup> |
| --- | --- | --- | --- | --- | --- | --- | --- | --- | --- |
|  |  |  | HACs vs DO | NACs vs DO | HACs vs NACs |  |  |  |  |
| PA1903 | phzE2 | phenazine biosynthesis protein PhzE | 30383.00 | --- | 30383.00 | Pyocyanin and phenazine biosynthesis (46,72) | 4 | 69.0 | 7.3 |
| PA2444 | glyA2 / ShrA | serine hydroxymethyltransferase | 21.96 | --- | --- | SCVs, biofilm formation, EPS production, motility, redox control (73) | 10 | 45.0 | 47.6 |
| PA3559 | PmrE | probable UDP-glucose dehydrogenase | 21.53 | --- | --- | EPS production, antibiotic resistance, LPS modification (74,75) | 3 | 51.8 | 10.6 |
| PA1327 | PA1327 | probable protease | 11.20 | --- | --- | Protease activity | 12 | 72.5 | 20.3 |
| PA3723 | PA3723 | probable FMN oxidoreductase | 5.46 | --- | --- | Stress resistance, antibiotic resistance (76) | 24 | 40.3 | 80.4 |
| PA4547 | PilR | two-component response regulator PilR | 4.52 | --- | --- | Motility, virulence (58) | 4 | 49.7 | 13.3 |
| PA4411 | MurC | UDP-N-acetylmuramate--alanine ligase | 3.05 | --- | --- | Antibiotic resistance, peptidoglycan biosynthesis (77,78) | 13 | 52.0 | 25.8 |
| PA1772 | RraA | probable methyltransferase. Ribonuclease activity regulator protein | 2.64 | --- | --- | RNA turnover, global regulator (79) | 11 | 17.4 | 67.3 |
| PA0083 | TssB1 | TssB1 | 2.46 | --- | --- | Antibiotic resistance, T6SS (80) | 13 | 18.8 | 73.8 |
| PA1178 | OprH | PhoP/Q and low Mg<sup>2+</sup> inducible outer membrane protein H1 precursor | 2.17 | --- | --- | Antibiotic resistance (81) | 23 | 21.6 | 69 |
| PA4964 | ParC | Topoisomerase IV subunit A | 1.77 | --- | --- | Antibiotic resistance (82,83) | 18 | 83.4 | 34.4 |
| PA0141 | Ppk2 | conserved hypothetical protein | --- | --- | 1.88 | Antibiotic resistance (84) | 20 | 40.8 | 72.8 |
| PA3262 | PA3262 | probable peptidyl-prolyl cis-trans isomerase, FkbP-type | --- | --- | 1.74 | Virulence; peptidoglycan biosynthesis (85) | 21 | 26.8 | 48.6 |
| PA4542 | ClpB | ClpB protein | --- | --- | 1.37 | Stress resistance, virulence (86) | 72 | 95.0 | 70.3 |
| PA4163 | Tse8 | hypothetical protein | -2.29 | --- | --- | Virulence (87) | 20 | 60.5 | 49.6 |
| PA0428 | RhlE2 | probable ATP-dependent RNA helicase | -2.38 | --- | --- | RNA turnover, global regulator, virulence (72) | 30 | 70.1 | 51 |
| PA2991 | Sth | soluble pyridine nucleotide transhydrogenase | -2.61 | --- | --- | Redox control (88) | 30 | 51.2 | 64.9 |
| PA2667 | MvaT/ MvaU | MvaU | -2.68 | --- | --- | Virulence, pyocyanin prod., motility, protease prod., biofilm formation, QS (89–91) | 19 | 13.4 | 87.2 |
| PA3083 | PepN | aminopeptidase N | -4.26 | -2.38 | --- | Virulence, competition (92) | 45 | 100.1 | 41.5 |
| PA5078 | OpgG | OpgG | -4.44 | -2.79 | --- | Stress resistance, biofilm formation (93) | 14 | 59.5 | 34.9 |
| PA3831 | PepA | leucine aminopeptidase | -5.07 | --- | --- | Virulence, protein turnover (94) | 33 | 52.3 | 71.1 |
| PA5507 | PA5508 | hypothetical protein | -5.56 | --- | --- | QS (95) | 12 | 24.3 | 68.2 |
| PA0301 | SpuE | polyamine transport protein | -6.02 | --- | --- | TSS3 regulation, virulence (96,97) | 12 | 40.1 | 55.9 |
| PA0423 | PasP | lipocalin protein family member | -6.04 | --- | --- | Antibiotic resistance, virulence (98,99) | 17 | 20.8 | 73.3 |
| PA1847 | NfuA | NfuA | -7.77 | --- | --- | Stress resistance, iron homeostasis, virulence (100,101) | 8 | 21.1 | 47.9 |
| PA3470 | PA3470 | NUDIX hydrolase | -105969.67 | --- | --- | Antibiotic & stress resistance, pyocyanin prod (102) | 3 | 16.7 | 13.2 |
| PA3115 | FimV | motility protein FimV | --- | --- | -1.40 | Motility, T2SS (103, 104) | 23 | 96.9 | 27.4 |
| PA4427 | SspB | stringent starvation protein B | --- | --- | -5.65 | Antibiotic resistance, virulence Iron homeostasis, QS (105) | 3 | 14.5 | 47.4 |
| PA4525 | PilA | type 4 fimbrial precursor PilA | --- | --- | -13.11 | Virulence, biofilm formation, motility (58,106) | 1 | 15.5 | 19.5 |
| PA2927 | PlaB | phospholipase B | --- | --- | -23843.67 | Biofilm formation (63) | 4 | 49.5 | 5.2 |

Blue shading represents proteins increased in abundance or unique to hypoxia. Dark blue: $\geq 5$ -fold increase in abundance; medium blue: 2- to 5-fold increase in abundance, light blue: 1.5- to 2-fold increase in abundance. Orange shading represents proteins decreased in abundance in hypoxia or absent in hypoxia. Dark orange: $\geq 5$ -fold decrease in abundance; medium orange: 2- to 5-fold decrease of abundance; light orange: 1.5- to 2-fold decrease in abundance.
<sup>a</sup>Gene ID – *P. aeruginosa* PAO1 gene locus ID.
<sup>b</sup>Proteins not significantly altered in abundance between given conditions are marked as “---”
<sup>c</sup>Relevant protein function assigned by literature searches, UniProt Database or *Pseudomonas* Genome Database.
<sup>d</sup>NP - Unique peptides: The number of peptide sequences unique to a protein
<sup>e</sup>MW - Molecular weight as determined by Q-Exactive LC-MS using *P. aeruginosa* PAO1 reference (kDa)
<sup>f</sup>SC - % sequence covered by matching peptides for the identified protein

We examined whether the alterations in protein abundance associated with antibiotic resistance affect affected antibiotic susceptibility of adapted cultures. Changes in sensitivity to a wide range of their antibiotic classes were observed in both HACs and NACs compared to the early infection strain (Fig 4A and 4B). However, both H1 and H2 populations clearly showed greater antibiotic resistance than NACs or ancestral cultures, correlating with the relative changes in protein abundance. Increased resistance to fluoroquinolones, macrolides, cephalosporins, carbapenems and penicillins was observed in both H1 and H2 cultures, together with decreased resistance to aztreonam. Interestingly, the H3 culture only showed an increased resistance to aztreonam with no change in susceptibility to any other antibiotic tested. All strains were resistant to tetracycline.

**Figure 4.**
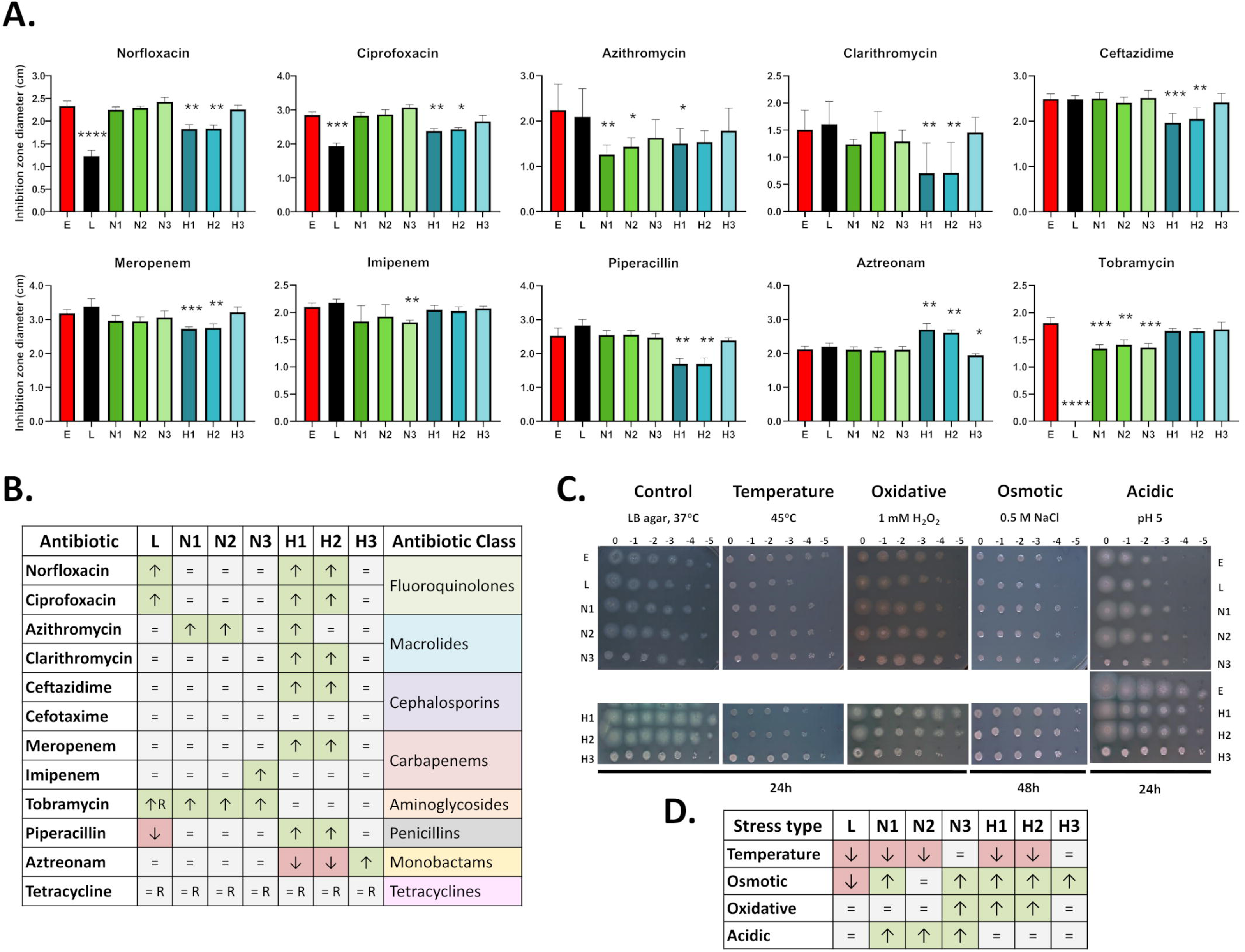
Effects of long-term hypoxia adaptation on antibiotic resistance and stress resistance. A. Graphs represent the antibiotic susceptibility of early infection control strain AMT 0023-30, late infection strain, AMT 0023-34, and 28-day adapted populations using diffusion discs, tested in normoxic conditions. Bars represent the mean inhibition zone diameter. The data represent three biological replicates and error bars represent the standard deviation. The statistical significance was determined with the nonparametric Kruskal-Wallis test in combination with uncorrected Dunns multiple comparisons test (*p< 0.05; **p< 0.01; ***p<0.001; ****p< 0.0001). B. Summary of antibiotic resistance to all tested antibiotics. C. Determination of stress resistance with droplet assays. Representative pictures of colonies from serial dilutions of each strain plated on agar with the specific stress factor added. The experiments were performed in normoxic conditions at least three times for each stress type with the early infection strain as a control. D. Table summarising changes in stress sensitivity. ↑ - increased resistance; ↓ - decreased resistance; = no significant difference in resistance; R – no inhibition zone observed. E: early infection clinical strain; L: late infection clinical strain; N1, N2, N3: 28-day normoxia-adapted populations; H1, H2, H3: 28-day hypoxia adapted populations.

### Hypoxia-adapted cultures show changes in stress response

Many proteins related to stress responses showed changes in abundance in response to long-term exposure to hypoxia (Table 4), including the osmotic stress response associated protein, OpgG, and the oxidative stress response associated protein, PA3723. The hypothetical protein, PA5139 previously associated with stress resistance, and lactate dehydrogenase LldD, associated with oxidative stress resistance were detected in the ancestral strain and HACs, but not any of the NACs.

To examine the impact of these alterations on the phenotype of each of the individual populations, droplet stress plates were prepared to test a range of stresses described as hallmarks of the CF lung environment. No clear pattern of resistance to high temperature, osmotic or acidic stress between the HACs and NACs was observed (Fig 4C and 4D). Interestingly, increased resistance to oxidative stress was observed in both H1 and H2 cultures and in the N3 culture, which were also the only populations where SCVs had been detected on day 28, and where increased biofilm production and exopolysaccharide production were observed.

### Hypoxia-adapted cultures show increased biofilm formation and exopolysaccharide production

Typically, anaerobic conditions promote biofilm formation, with greater numbers of live bacteria, increased thickness, and more compact structures (107). In this work, changes in the abundance of proteins associated with biofilm and/or exopolysaccharide production were also observed in response to exposure to 6% oxygen (Table 3 and Table 4). Two proteins associated with biofilm associated antibiotic resistance were increased in abundance, while three were decreased. Additionally, some of the proteins associated with biofilm formation or EPS production were not detected in at least one of the groups: exopolysaccharide export protein, PelC, was identified solely in HACs but not in NACs or ancestral strain (Table 3); solute-binding protein, PA5139, is present in the ancestral strain, maintained in HACs, while undetectable in NACs; the polymer-forming cytoskeletal protein PA0664 is detected in the ancestral strain, and in two HACs, but none of the NACs; phospholipase PlaB was detected in two replicates of the ancestral strain, detected in all of the NACs, but none of the HACs.

To confirm the impact of these proteome changes on biofilm and exopolysaccharide production, these phenotypes were examined in each individual adapted culture (Fig 5A and 5B). Both H1 and H3 populations showed a significant, dramatic increase in biofilm formation as compared to the early infection strain after 24 h (6.1- and 4.3-fold increases respectively). Significant increases in biofilm production were also observed in the late infection strain (1.6-fold), H1 (3.9-fold) and H2 (1.4-fold) over the ancestral strain at 48 h, and this trend persisted at 72h (Fig 5A). The increased biofilm production of the H3 population was transient and reverted to AMT 0023-30 early infection strain levels by 48 h. Congo Red staining revealed darker colonies for H1 and H2 populations which were more wrinkly than in the other strains indicating a higher production of exopolysaccharides (Fig 5B). Interestingly, a similar phenotype was observed in culture N3, again suggesting that this phenotype is not solely dependent on oxygen availability.

**Figure 5.**
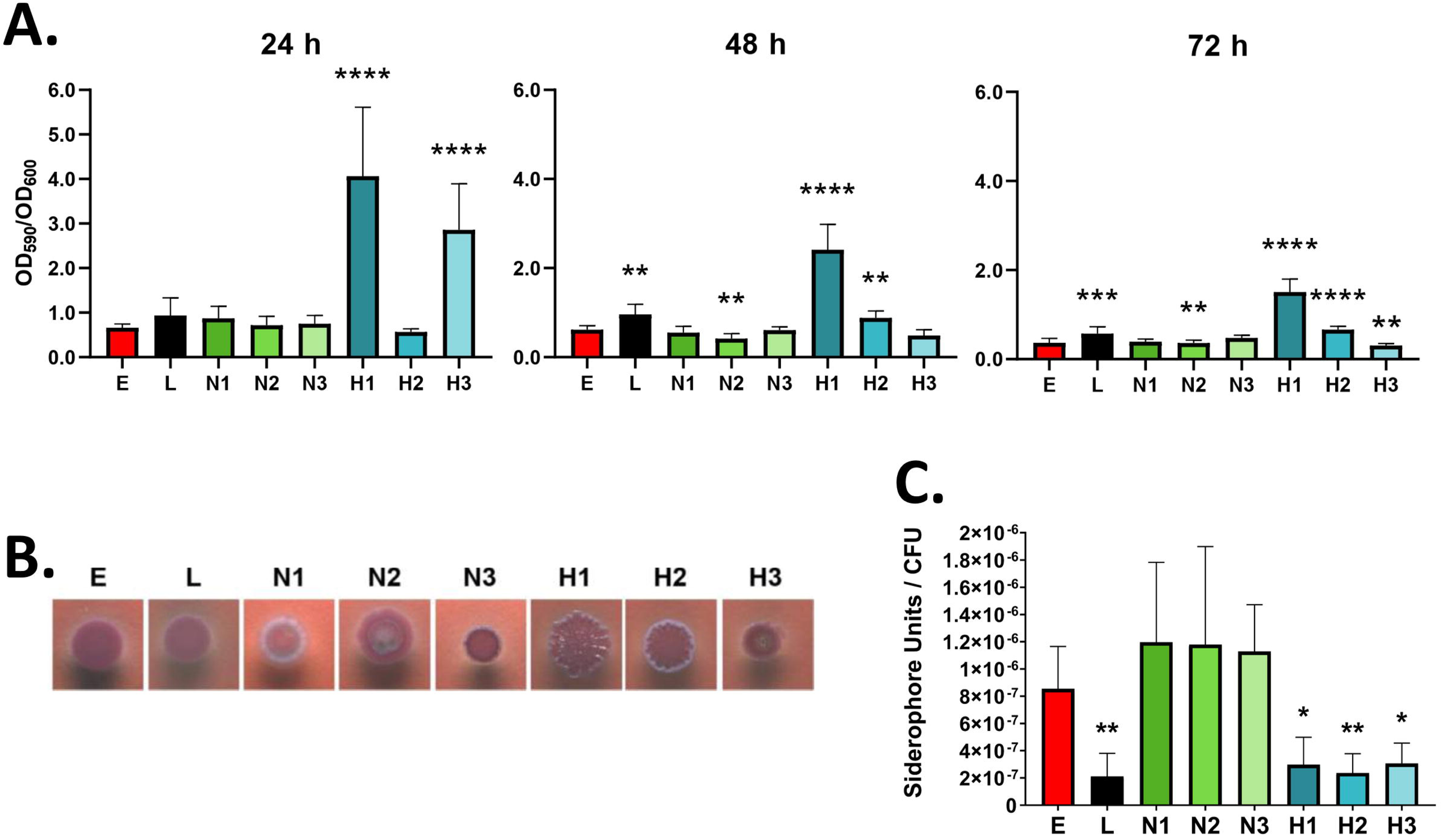
Effects of long-term hypoxia adaptation on biofilm formation, exopolysaccharide production and siderophore production. A. Biofilm formation capability of hypoxia-adapted cultures determined with the crystal violet assay after 24, 48 or 72h of static incubation on a 96-well plate in normoxic conditions. Data represent the mean value of the OD_590_/OD_600_ ratio from three separate experiments with at least six technical replicates for each strain. Statistically significant difference by Kruskal-Wallis one-way ANOVA test compared to the ancestral, early infection strain (E) with an uncorrected Dunn’s test (**p< 0.01; ***p<0.001; ****p< 0.0001). B. Exopolysaccharide production by the adapted cultures. The photographs show colonies of different populations of bacteria. The change of colour to a darker shade and a wrinkly colony surface indicate a higher exopolysaccharide production. C. Siderophore production represented as the mean of siderophore units produced by each colony forming unit, tested in normoxic conditions. Statistically significant differences by ordinary one-way ANOVA test compared to the ancestral, early infection strain with an uncorrected Fishers LSD test are shown (*p<0.05; **p< 0.01). E: early infection clinical strain (AMT 0023-30); L: late infection clinical strain (AMT 0023-34); N1, N2, N3: 28-day normoxia-adapted populations; H1, H2, H3: 28-day hypoxia adapted populations.

Figure 5. Effects of long-term hypoxia adaptation on biofilm formation, exopolysaccharide production and siderophore production. A. Biofilm formation capability of hypoxia-adapted cultures determined with the crystal violet assay after 24, 48 or 72h of static incubation on a 96-well plate in normoxic conditions. Data represent the mean value of the OD_590_/OD_600_ ratio from three separate experiments with at least six technical replicates for each strain. Statistically significant difference by Kruskal-Wallis one-way ANOVA test compared to the ancestral, early infection strain (E) with an uncorrected Dunn’s test (**p< 0.01; ***p<0.001; ****p< 0.0001). B. Exopolysaccharide production by the adapted cultures. The photographs show colonies of different populations of bacteria. The change of colour to a darker shade and a wrinkly colony surface indicate a higher exopolysaccharide production. C. Siderophore production represented as the mean of siderophore units produced by each colony forming unit, tested in normoxic conditions. Statistically significant differences by ordinary one-way ANOVA test compared to the ancestral, early infection strain with an uncorrected Fishers LSD test are shown (*p<0.05; **p< 0.01). E: early infection clinical strain (AMT 0023-30); L: late infection clinical strain (AMT 0023-34); N1, N2, N3: 28-day normoxia-adapted populations; H1, H2, H3: 28-day hypoxia adapted populations.

### Adaptation to hypoxia affected iron acquisition and homeostasis

Proteomic analysis showed changes in the abundance of proteins related to iron homeostasis and acquisition pathways (Table 4). CyaY protein involved in iron-sulphur cluster assembly was undetected in HACs but was present in the ancestral strain and maintained in the NACs (Table 3). NfuA, previously associated with iron homeostasis was decreased in abundance by nearly 8-fold relative to the ancestral strain (Table 4). In contrast, the stringent starvation protein SspB was increased in abundance over 5-fold in HACs compared to NACs.

Differences in iron acquisition and siderophore levels of all individual populations showed a consistent, clear decrease in siderophore production in all HACs by 2.8- to 3.6-fold but no significant change in NACs (Fig 5D). No significant differences were detected in the pyochelin and pyoverdine production pathways, but the PchR transcriptional regulator was undetectable in all the replicates of the ancestral strain, two HACs and present in all NACs. Moreover, a 4-fold decrease in siderophores was also observed between the early infection strain (ancestral) and late infection strain.

### Adaptation to hypoxia induces changes in motility

A common adaptation associated with P. aeruginosa chronic infection isolates is loss of motility. Multiple proteins associated with motility showed changes in abundance in the HACs. Flagellar hook protein FlgE, and spermidine dehydrogenase SpdH were undetectable in the HACs while present in the ancestral strains and NACs (Table 3). The two-component response regulator, PilR, and serine hydroxymethyltransferase ShrA were increased in abundance in HACs (4.5 and 22.0-fold respectively), but not in NACs compared to the ancestral strain. In contrast, DNA-binding protein MvaT was decreased in abundance by 2.7-fold. Additionally, type 4 fimbrial precursor PilA, and FimV were both decreased in abundance (13- and 1.4 fold respectively) in HACs vs NACs. Notably, many of these proteins are also associated with biofilm formation and/or antibiotic resistance (ShrA, MvaT, PilA, FlgE).

To elucidate the effect of hypoxia on motility further, the motility of the individual cultures was assessed. The late infection strain showed a 2-fold reduction in swarming (Fig 6B) and a moderate reduction in twitching motility (1.3-fold) (Fig 6C), while H1 and H2 showed an increased swimming (1.5 and 1.2-fold) compared to the early strain (Fig 6A). All forms of motility were decreased in H3 by over 2.4-fold (Fig 6A, 6B, 6C). A similar pattern was observed in culture N3, with a 1.6-fold decrease in swimming motility and a 1.9-fold decrease in twitching motility (Fig 6A, 6C). The pattern of swarming motility was clearly different between NACs and HACs. N1 and N2 populations showed classic swarming patterns with clearly distinguishable dendrites despite showing no changes in diameter (Fig 6B). N3 showed a different pattern, forming colonies with smooth edges. The H1 and H2 colonies on swarm plates were clearly distinguishable from the N1 and N2 counterparts, with no clear dendrites being formed (Fig 6). Overall, exposure to hypoxia led to clear changes in strain motility.

**Figure 6.**
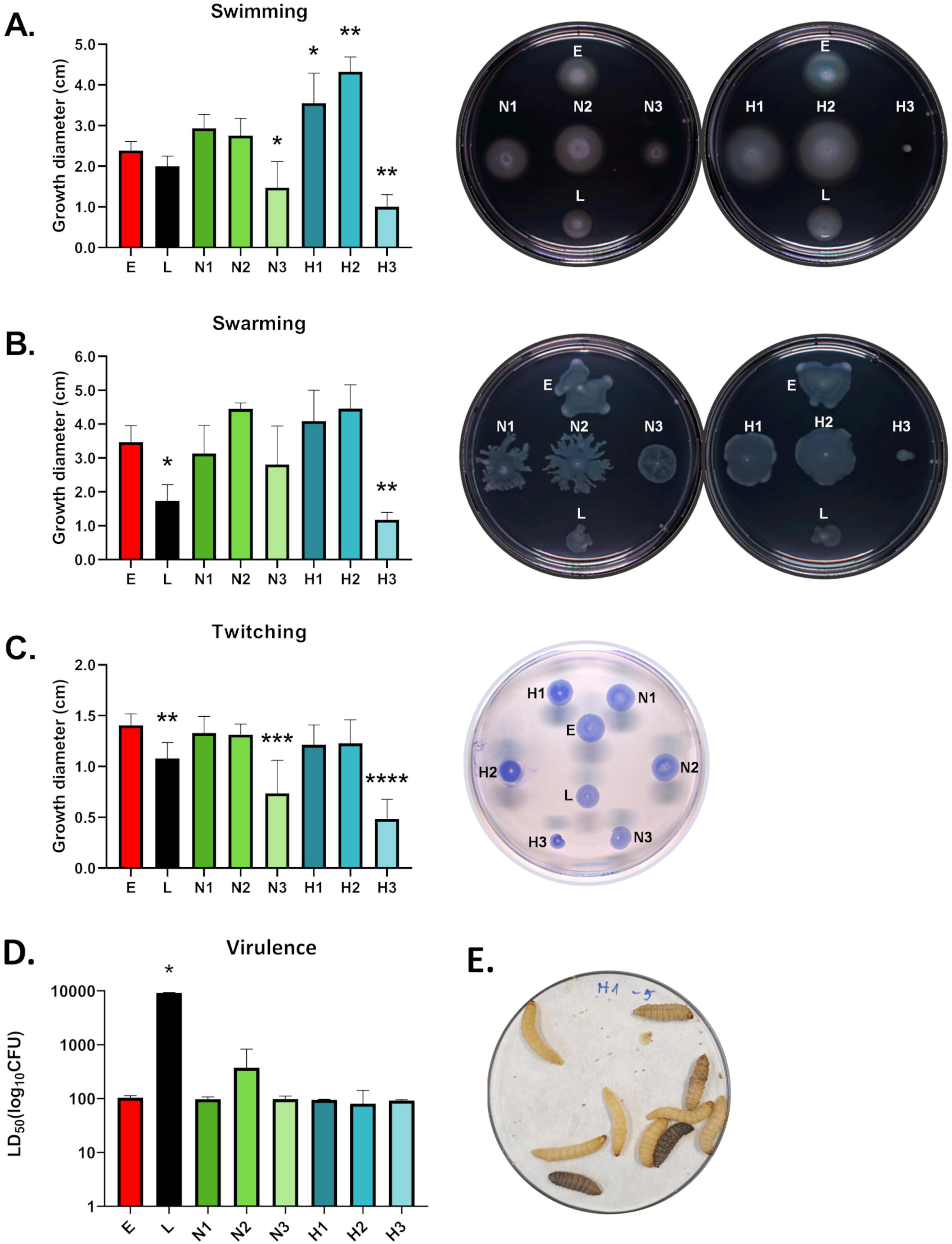
Effects of long-term hypoxia adaptation on P. aeruginosa motility and virulence in the Galleria mellonella model. A. Changes in swimming motility. B. Changes in swarming motility. C. Changes in twitching motility. For each type of motility, a graph representing the mean of diameter of the colony with bars representing data from at least three separate experiments conducted in normoxic conditions, with at least two technical replicates. Statistically significant difference by Kruskal-Wallis one-way ANOVA test compared to the ancestral, early infection strain (E) with an uncorrected Dunn’s test (*p< 0.05; **p< 0.01; ***p<0.001; ****p< 0.0001). D. Graph representing the mean CFU required to reach a lethal dose leading to 50% mortality (LD50) within 24h of the experiment from at least three independent experiments. A touch to the larvae head was used to determine if they are alive. Statistically significant difference by Kruskal-Wallis one-way ANOVA test compared to the ancestral, early infection strain (E) with an uncorrected Dunn’s test is presented (*p< 0.05). E. A representative image showing the G. mellonella larvae melanisation. Healthy larvae with no disease symptoms are a bright cream colour. Larvae showing a slightly beige colour show first symptoms of infection, larvae with dark spots/stipes are clearly affected and the severely melanised larvae are in the late stages of the infection or dead. E: early infection clinical strain AMT 0023-30; L: late infection clinical strain AMT 0023-34; N1, N2, N3: 28-day normoxia-adapted populations; H1, H2, H3: 28 day hypoxia adapted populations.

### Effects of hypoxia-adaptation on in vivo virulence

Previously we showed that three different late infection CF isolates were less virulent in the *G. mellonella* in vivo model than their earlier infection counterparts (37). HACs showed that adaptation to hypoxia altered the abundance of several proteins involved in virulence, and many virulence-associated phenotypes, including SCV formation, stress and antibiotic resistance, toxin and LPS production, and motility. As mentioned, phenazine biosynthesis protein PhzE was unique to HACs; Iron(II)/α-ketoglutarate-dependent oxygenase AmbC involved in toxin production was undetectable in the ancestral strain and two of the NACs while present in all HACs. The earlier mentioned two component system proteins GacS and ColR were undetectable in all HACs while present in two replicates of the ancestral strain and all NACs (Table 3). Additionally, virulence factors, leucine aminopeptidase PepA and lipocalin protein PasP, were decreased in abundance in HACs (Table 4). Unexpectedly, no differences in virulence were observed *in vivo* in *G. mellonella* between the ancestral, early infection strain and the HACs (Fig. 6D and 6E), showing hypoxia is unlikely to drive this adaptation.

## Discussion

Understanding the complex network of pathway interactions and the evolutionary pressures *P. aeruginosa* undergoes in different environments is the key to understanding its biology and the mechanisms behind chronic infection, thereby, facilitating the development of effective therapeutics. Hypoxic niches are abundant in the CF lung due to increased mucus thickness, obstructive mucus plugging and increased oxygen consumption by epithelial and immune cells (28,108,109). Studies have shown that microaerobic respiration is the predominant mode of *P. aeruginosa* growth in the CF lung (110,111), with two transcription factors, Anr and Dnr, tightly regulating the response to changes in oxygen tension (29,71,112,113).

While the impact of short-term severe hypoxia and anoxia on *P. aeruginosa* has been previously described (114,115), the impact of moderately-low hypoxia stress has not been examined over extended periods. To enable an in-depth phenotype and proteome analysis of adapted cultures, we restricted the study to three replicates in each condition. This focused approach allowed us to demonstrate that prolonged low oxygen conditions alone can induce stable changes to the proteome and phenotype, many of which are typically associated with chronic infection, and thus may facilitate the transition from acute to chronic infection.

### Diverse patterns of adaptation to long-term hypoxia

We observed two distinct patterns of hypoxia adaptation in *P. aeruginosa*, with H1 and H2 adapting differently from H3 (Fig. 7). This is consistent with reports on evolution of different characteristics under the same environmental pressures, including bacterial populations isolated from the same lung niche, or in different regions of a biofilm, and enables bacterial competitiveness (21,116–120). Flynn et. al (2016), using a similar experimental evolution approach compared three biofilm cultures and three planktonic cultures of *P. aeruginosa* PA14 over 90 days and determined that diverse phenotypes appeared over time with different abilities to persist between replicate cultures, and also between colonies isolated from the same population (117). Furthermore, the various colony morphologies observed in their study correlated with temporal variations in c-di-GMP levels (117), which aligns with the changes in the abundance of proteins associated with the response to c-di-GMP in HACs. We will evaluate the potential genetic changes that may have led to this in a future study.

**Figure 7.**
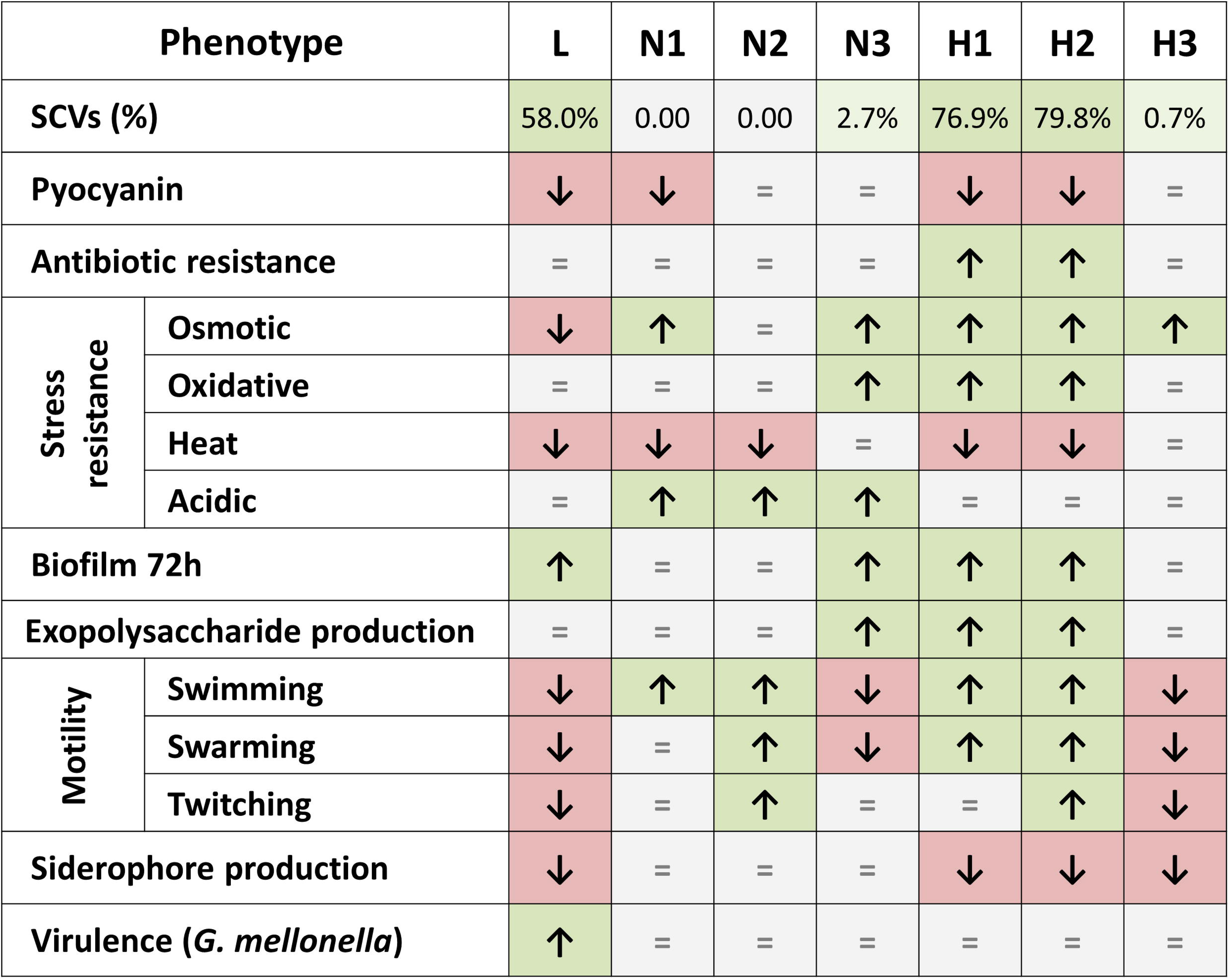
Patterns of adaptation to long-term hypoxia. Summary of the phenotypic changes of adapted cultures tested in normoxic conditions compared to the ancestral strain AMT 0023-30 (E) and the late infection strain AMT 0023-34 (L). It should be noted that L is a single clinical strain and not representative of all chronic infection isolates or strains. ↑ - indicates an increase; ↓ - indicates a decrease; = indicates no significant difference in resistance; L: late infection clinical isolate; N1, N2, N3: 28-day normoxia-adapted populations; H1, H2, H3: 28-day hypoxia adapted populations. (121).

Two of the HACs in this study (H1 and H2) showed consistent alterations across multiple phenotypes, while the third population displayed only slight alterations relative to the ancestral, early infection strain and was closer in phenotype to the NACs (Fig. 7). This pattern aligns with earlier observations that proteome changes caused by short term hypoxia response were strain-specific (115), suggesting that different adaptation patterns can occur as the population diversifies over the longer term. Given the diversity of *P. aeruginosa* isolates within patients and between patients, it should be noted that the late infection strain which was examined in parallel in phenotyping experiments represents one branch within a potentially more complex evolutionary tree in that patient. Further genetic characterisation of the populations and specific subpopulations will provide additional information on diversity.

### Adaptation of P. aeruginosa to long-term hypoxia is consistent with changes observed during chronic infection

When oxygen availability decreases, electron transport chain efficiency also decreases, leading to NADH accumulation and an increased NADH/NAD⁺ ratio (122), creating a more reducing intracellular environment (defined as redox stress) which must be mitigated to maintain cellular function and viability. This is regulated by c-di-GMP signalling, which plays a central role in mediating low oxygen responses and stress resistance in *P. aeruginosa* (123,124). It facilitates metabolic switching in response to environmental stimuli, including the transition from a planktonic to a biofilm lifestyle (24,124,125) and regulates multiple phenotypes such as EPS production, motility, pyocyanin production, and antibiotic resistance (126–128).

Based on previous studies (113,115,129,71), we expected decreased oxygen availability would lead to an increased abundance of the Anr regulator. However, in contrast, Anr was not detected in any of the HACs, although it was present in the ancestral strain. Previous studies investigating the Anr regulon were performed under either anoxia or severe hypoxia (0.2-2% O_2_), rather than the moderately-low hypoxia used here. There are three possible explanations: 1) the oxygen level may be too high for Anr activity; 2) there are other levels of regulation of Anr as yet undiscovered; 3) hypoxia adaptation may have selected for strains that downregulate Anr to reduce energy costs. This is important in the context of the CF lungs, where oxygen gradients from anoxia to physiological oxygen are present, suggesting more variable responses and adaptation mechanisms than assumed in previous studies.

Given the absence of Anr in HACs, alternative regulatory pathways are likely, as exemplified by >20-fold increase in abundance of the serine hydroxy methyltransferase, ShrA in HACs. Deletion of *shrA* was previously shown to lead to increased c-di-GMP levels, emergence of SCVs, increased biofilm, EPS and pyocyanin production, decreased swarming motility and suppression of iron acquisition genes and denitrification operons (Fig 8A) (73) which align with many of the phenotypes identified in response to long-term hypoxia. We propose that the redox imbalance caused by hypoxia increases the abundance of ShrA *via* a yet unknown mechanism.

**Figure 8.**
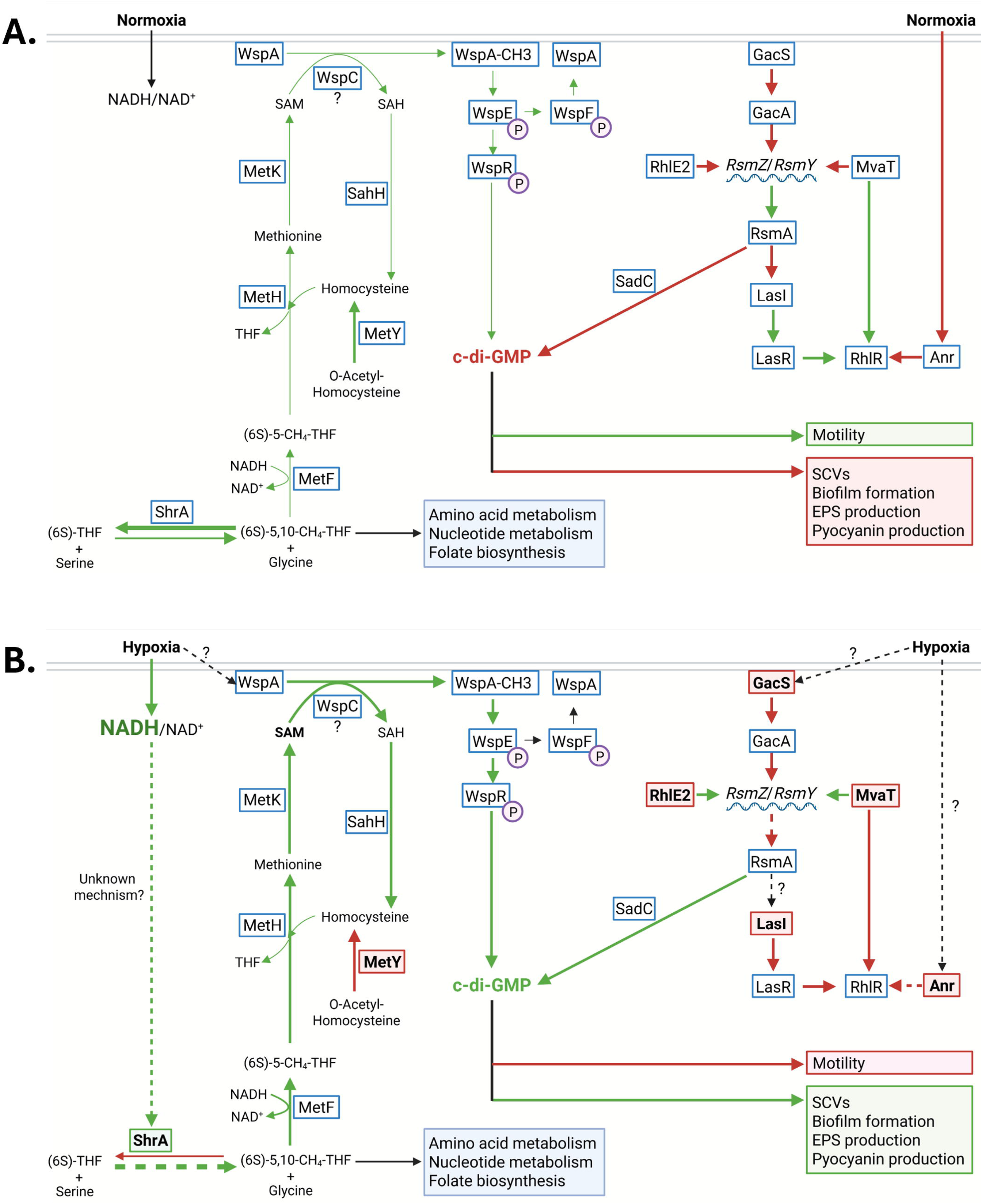
Suggested mechanism of regulation of c-di-GMP levels by 6% oxygen hypoxia exposure. A. Mechanism of regulation of c-di-GMP levels by ShrA and the Gac/Rsm system system in normoxic conditions based on available literature. When NADH/NAD+ ratio is balanced, ShrA activity leads mostly to the production of (6S)-THF, which leads to limited SAM production and WspA methylation. This leads to a low production of c-di-GMP by WspR. B. Proposed mechanism of regulation of c-di-GMP levels by ShrA and the Gac/Rsm system in hypoxic conditions based on the proteomic analysis and phenotype studies presented in this work. An increased NADH/NAD+ ratio may lead to a shift ShrA activity via an yet unknown mechanism, which leads to an increased (6S)-5,10-CH₄-THF production, increasing SAM production and WspA methylation. This leads to a increased production of c-di-GMP by WspR. Anr: anaerobic regulator; GacA: response regulator; GacS: sensor kinase; LasI: acyl-homoserine lactone (AHL) synthase; LasR: Las quorum sensing system transcriptional regulator; MetF: methylenetetrahydrofolate reductase; MetH: methionine synthase; MetK: S-adenosylmethionine synthetase; MetY: homocysteine synthase; MvaT: hitone-like nucleoid structuring global transcriptional regulator; RhlE2: Dead-boxRNA helicase; RsmA: RNA-binding regulator of secondary metabolites A; RsmY: rsm system small regulatory RNA Y; RsmZ: rsm system small regulatory RNA Z; SadC: suppressor of alginate production diguanylate cyclase; ShrA: serine hydroxymethyltransferase; WspA: methyl-accepting chemotaxis protein; WspC: methyltransferase; WspE: histidine kinase; WspF: methylesterase; WspR: diguanylate cyclase; THF: tetrahydrofolate; (6S)-THF: (6S)-tetrahydrofolate; (6S)-5,10-CH₄-THF: (6S)-5,10-methylenetetrahydrofolate; SAM: S-adenosylmethionine; SAH: S-adenosylhomocysteine. Green arrows indicate upregulation or activation. Red arrows indicate downregulation or inactivation. Solid lines define literature described relationships. Dashed lines define hypothetical relationships. Green boxes indicate proteins increased in abundance in the proteomic analysis performed as a part of this work. Red boxes indicate proteins decreased in abundance in the proteomic analysis performed as a part of this work.

ShrA catalyses the reversible conversion of glycine and (6S)-5,10-methylene-THF ((6S)-5,10-CH₄-THF) to serine and (6S)-THF, with a nearly threefold preference for the latter products (73). Besides its role in c-di-GMP biosynthesis regulation, 5,10-CH_4_-THF regulates folate and nucleotide biosynthesis and amino acid metabolism. However, as these findings were identified under normoxic conditions, ShrA’s role in hypoxia remains unclear and warrants further investigation. In particular, its reaction flux may be influenced by factors such as the increased NADH/NAD⁺ ratio observed under hypoxic conditions. Given that ShrA contributes to restoring redox balance by reducing the NADH/NAD⁺ ratio, its elevated abundance is likely to be beneficial under hypoxic stress. The (6S)-5-CH₃-THF synthesized by ShrA is further metabolized through several steps to produce S-adenosylmethionine (SAM), which serves as methyl group donor for the methylation of WspA (73). Methylated WspA activates WspR and c-di-GMP synthesis (130). Thus, a shift towards synthesis of (6S)-5-CH_4_-THF would increase the methylation of WspA, leading to increased levels of c-di-GMP under hypoxic conditions (Fig 8B). We acknowledge that ShrA regulation in hypoxia may be more complex and requires further investigation, as it has been previously shown to be induced by homocysteine, repressed by SAM, and activated via quorum sensing and PpyR (131,132).

Sensory histidine kinase GacS is also involved in c-di-GMP regulation. Under non-stress conditions, GacS levels are naturally low or undetectable (61), however, when GacS is absent, less GacA is phosphorylated, leading to reduced expression of RsmY and RsmZ sRNAs (133). This results in a larger pool of active RsmA, increased repression of target mRNAs, and ultimately, lower c-di-GMP levels. Unexpectedly, we detected GacS in the ancestral strain and NACs, but not the HACs which is possibly due to the activity of regulators of RsmZ/RsmY sRNAs. Histone-like protein MvaT reduces c-di-GMP levels by binding to the upstream regions of rsmY/rsmZ repressing their transcription, increasing RsmA levels and repressing the activity of diguanylate cyclases (90). Additionally, the RhlE2 reduces RsmY and RsmZ stability, thus promoting RsmA sequestering and allowing the production of c-di-GMP (72). In HACs we observed comparable >2.5-fold decreases in the abundance of MvaT, and RhlE2 compared to the ancestral strain, which was not observed in NACs. Therefore, we speculate that the presence of GacS in NACs and ancestral strain is overcome by the activity of RhlE2 and MvaT coupled with the decreased production of c-di-GMP by WspR.

Taken together, our findings suggested that the observed changes in abundance of ShrA, GacS, RhlE2 and MvaT lead to an increase in cellular c-di-GMP levels, consistent with the observed phenotypes including increased biofilm and EPS production, emergence of stable SCV phenotypes, alterations in motility, antibiotic resistance and pyocyanin production which could also be associated with this signalling molecule. Thus, we hypothesised that while Anr primarily responds under anoxic conditions or severe hypoxia, other proteins, including ShrA, may regulate relevant pathways under moderately-low O_2_ levels after long-term exposure to hypoxia. However, we have not observed any changes in the intracellular concentrations of c-di-GMP levels between the ancestral early infection *P. aeruginosa* strain and any of the HACs and NACs or the late infection strain despite the observed changes in related phenotypes (S6. Figure). This was not unexpected as Yan et al. (2017) previously investigated the correlation between c-di-GMP levels, motility, and biofilm formation (134) and did not observe any direct correlation, despite using well-established models describing the mechanisms of c-di-GMP action. This finding suggests a more complex regulatory network, where the final phenotypic outcome depends on the cumulative effect of multiple small-scale changes rather than a straightforward 1:1 correlation. Such complexity could potentially explain the seemingly contradictory observations in c-di-GMP concentrations, regulator protein abundances, and variation in motility and biofilm formation.

Interestingly, the adaptation of *P. aeruginosa* to long-term hypoxia mirrors the phenotype and molecular changes previously reported for *P. aeruginosa* chronic infection isolates (8,18,21,23,36,65,68), including increased SCV prevalence, increased biofilm and exopolysaccharide production, increased antibiotic resistance and decreased siderophore production. Importantly, all these changes are stable, because all phenotypes were examined in normoxic conditions; thus, they are a result of adaptation to hypoxia, rather than a plastic temporary response to hypoxic conditions.

While a potential limitation of our experiment is that growth in relatively rich laboratory media does not represent the conditions in the lung, the fact that we can observe changes resembling adaptation to chronic infection even in the absence of nutrient stress, highlights a clear role for hypoxia in this process. Choosing a planktonic model for this study allowed us to balance the feasibility of the experimental design and the relevance of the model in the clinical context but considering the obtained results we argue that this approach effectively reveals the potential role of long-term hypoxia in driving adaptation during chronic infection.

Two distinct SCV types emerged in the long term cultures which were more prevalent in the HACs, and one of the morphotypes was exclusive to hypoxia. We observed a temporary emergence of SCV 1 in the NACs, which was not positively selected in normoxia, possibly due to the lack of an adaptive benefit long-term. Under optimal laboratory conditions, long-term cultivation can induce stress due to factors such as high population density, accumulation of metabolic byproducts, nutrient gradients leading to transient SCV arising in the population. *P. aeruginosa* also possesses phase variation systems that allow spontaneous phenotypic switching, generating SCVs even in the absence of external stressors (135,136). These variants may be maintained at low frequencies, preparing the population for potential environmental shifts (135). The abundance of c-di-GMP signalling pathways proteins, Wsp, and YfiBNR, typically associated with SCVs (41,135), was unchanged in the HACs, but as mentioned earlier, the ShrA protein indirectly regulates the Wsp system activity. Additionally, the inactivation of *gacS* has been shown to be associated with the emergence of SCVs under stress conditions and suppression of the reversion of this phenotype to a normal colony variant (62). The latter may explain why the SCV populations in the NACs were transient, as GacS is expressed in the NACs but not HACs. SCVs are strongly associated with pathogenicity and persistence among several bacterial pathogens (137–139), and are prevalent among chronic *P. aeruginosa* isolates (137,138,140). Specifically, SCVs of *P. aeruginosa* correlate with increased exopolysaccharide and biofilm production, antimicrobial resistance, stress resistance and host immune responses (41,140). Their emergence has been linked to redox imbalances which can be caused by low oxygen conditions (136), although other environmental stressors or co-colonisation with other species also contribute to their increased prevalence (137,140,141). This confirms directly, for the first time, that prolonged hypoxia exposure is associated with increased SCV occurrence in *P. aeruginosa* populations. Identification of the factors that contribute to SCV emergence is crucial due to the challenges they pose in clinical settings (137) and recognising the environmental factors that select this phenotype may help us limit their occurrence. Both HACs that showed a high SCV prevalence (H1, H2) also showed increased resistance to a broad spectrum of antimicrobials together with increased biofilm and exopolysaccharide production. Surprisingly, no trends in motility relating to hypoxia exposure or biofilm formation were observed, although some changes in patterns of swarming colonies were noted. Further investigation of NCVs and SCV 1 and SCV 2 (genotypes and phenotypes) will provide insight on the contribution of the specific subpopulations to the observed changes in other adaptations and are ongoing.

Chronic infection *P. aeruginosa* CF isolates typically exhibit increased biofilm and exopolysaccharide production, and reduced motility (18,120,142–144), with biofilm formation relying on hypoxia responses due to the presence of steep oxygen gradients (30,145,146). Moreover, the awareness of the role of biofilms in antibiotic treatment efficiency is being increasingly acknowledged (147,148). Proteins associated with biofilm formation, exopolysaccharide production, and motility were changed in abundance. Notably, outer membrane lipoprotein PelC, which is essential for biofilm formation (45), was unique to HACs while in contrast, the flagellar hook protein FlgE, typically associated with motility, host interactions and immune evasion (54), was absent in HACs. Both are regulated by FleQ, a c-di-GMP binding transcriptional regulator, which activates the *pel* operon and inhibits the flagellar gene expression, coordinating this lifestyle switch (149). Two component system regulator PilR associated with increased biofilm formation, host cell attachment, and motility (58) was increased in abundance in HACs. However, the PilR-activated type IV fimbrial precursor, PilA, was decreased in abundance by >13-fold in HACs compared to NACs which suggests alternative regulation mechanisms are at play. Further highlighting this complexity, phospholipase PlaB, showing activity towards exogenous lipids in the biofilm matrix (63), was absent in HACs while UDP-glucose dehydrogenase PA3559, part of the EPS biosynthesis pathway (74), was increased in abundance in HACs. These changes collectively indicate an adaptive shift towards a biofilm lifestyle in HACs, aligning with changes observed in chronic infection adaptation.

Antibiotic resistance is a major hallmark of *P. aeruginosa* infections in CF (150), and short-term hypoxic and anaerobic conditions have been previously shown to increase *P. aeruginosa* antibiotic resistance (151–153). We demonstrate that long-term exposure to hypoxia induces stable changes in antibiotic resistance, which importantly, didn’t revert during in normoxic conditions used for phenotyping. Two of the HACs showed increased resistance to a wide range of antibiotics from different classes, further supporting the evidence of the extent of the influence of hypoxia pre-adaptation on drug resistance and the wide range of diverse mechanisms affected. Interestingly, high c-di-GMP levels have been shown to corelate with increased resistance to antibiotic, independently form the biofilm/ planktonic lifestyle (128). Several proteins involved in lipid A/LPS biosynthesis (LpxA, LpxB, WaaG), membrane stability (DsbG, ColR, OprH), and peptidoglycan biosynthesis (MurC, PA3262), many of which have been previously linked to antibiotic resistance (154–158) were altered in abundance. These may contribute to the observed broad-spectrum antibiotic resistance by affecting cell envelope permeability and structure. Moreover, other proteins linked to antibiotic resistance including phosphate kinase Ppk2, previously linked to multidrug resistance (84), and DNA topoisomerase IV subunit ParC involved in fluoroquinolone resistance (83,159) were increased in HACs (106).

The airways of people with CF are reported to show an increased abundance of available iron (160,161). *P. aeruginosa* expresses many iron acquisition systems including siderophore production, haem acquisition systems and complexes importing iron-bound citrate or catechol (162). Importantly, a strong positive correlation between increased sputum iron and chronic *P. aeruginosa* infections has been shown (163). Furthermore, iron uptake and homeostasis are regulated by the c-di-GMP through the Gac/Rsm cascade (164), in line with other alterations observed and underlines the role of hypoxia leading to changes in iron acquisition on chronic infection development, which led to recognising them as therapy targets for *P. aeruginosa* treatment (161,165). Surprisingly, we didn’t observe any alterations associated with pyochelin or pyoverdine biosynthesis and transport, although a clear decrease in siderophore production was observed in HACs. Notably, the PchR regulator was not detected in the ancestral strain, nor in two of the HACs, while present in all of the NACs, suggesting alternative mechanisms of control of siderophore production or transport. Iron-sulphur cluster proteins are essential for maintaining iron homeostasis and regulating diverse cellular functions (166). Notably, two iron-cluster proteins frataxin-like protein CyaY, and iron-sulphur cluster scaffold protein NfuA were absent or decreased in abundance in HACs, respectively. Additionally, the RhlE2 RNA helicase mentioned earlier has been shown to regulate expression of iron acquisition genes. These changes may explain the decreased siderophore levels observed, but require further investigation; notably, they reflect the downregulation of siderophore production by chronic infection isolates from CF lungs (167).

Adaptation of *P. aeruginosa* to chronic infection is frequently associated with reduction in expression of multiple virulence factors including pyocyanin, proteases, lipases, urease, elastase and secretion systems (18). Moreover, exposure to hypoxia attenuated the virulence of isolates from acute, but not chronic infections (168) and decreased their interactions with host cells, demonstrating the importance of this environmental pressure. Likewise, proteomic analysis of HACs, showed a decrease in abundance of many virulence factors: proteases (PasP serine protease, aminopeptidase PepA, aminopeptidase PepN), Phospholipase B, and type VI secretion system (TssB1, Tse8 effector protein) and type II secretion system effector proteins (PasP, PilA). LasI, a component of the QS system was not detected in HACs or the ancestral strain, but present in NACs. Since it positively regulates protease and lipase synthesis (and indirectly T6SS), its decreased abundance is consistent with the other alterations. Interestingly, phenazine biosynthesis protein PhzE was unique to HACs, which would suggest an increase in pyocyanin production in all HACs, however, we detected a change in pyocyanin in only two cultures. Thus, we might have expected decreased virulence in HACs, but did not observe any change in the *G. mellonella* infection model following hypoxia exposure. However, this acute infection model may not be optimal to assess the impact of long-term hypoxia adaptation on chronic infection.

Some of the observed changes in phenotypes, including increased biofilm and EPS production, and antibiotic resistance may result from the increased prevalence of the SCVs in the population, as all of these are altered in HAC 1 and 2, while limited changes were observed in HAC 3. Investigation of this is ongoing, however, given that many alterations in the proteome were conserved across all three HACs, despite divergent phenotypic outcomes suggests a more complex explanation involving additional, SCV-independent adaptive mechanisms.

A recent study defined *P. aeruginosa* 224 genes as pathoadaptive, and demonstrated that specific gene expression was associated with a host preference towards people with CF (169), many of which encode proteins altered in response to long-term hypoxia. Three of these, ColR, GacS and MexS were not detected in HACs, while another, PA0664 protein was maintained in HACs while not detected in NACs. Additionally, PilR and ParC were increased in abundance in HACs respectively, while OprM was decreased in abundance. Moreover, overall changes in the abundance of 15 proteins encoded by genes that were associated with CF host preference (including *oprM*, *pepA*, *fimV*, and *pilR*) and 11 that were associated with non-CF infections (including *mvaT* and *tssB1*) (169), suggest long-term hypoxia may play a role in *P. aeruginosa* infection both in CF and non-CF cohorts.

### Common adaptation mechanisms of opportunistic pathogens to hypoxia

Several oxygen-sensing mechanisms, homologous genes in hypoxia regulons and several phenotypes altered in chronic infection are shared across other opportunistic bacteria causing pulmonary infections. This suggests that adaptation to prolonged hypoxia could be a shared mechanism between many low oxygen and/or microaerophilic habitats and not limited to human infections. These environments are increasingly abundant in water and soil, where opportunistic bacteria reservoirs exist, and play significant roles in shaping their physiology, contributing to their increased prevalence (170–174). As many opportunistic bacterial pathogens causing CF infections reside in these niches, common adaptations are not unexpected.

Proteome analysis of HACs showed an increase in abundance of only two proteins encoded on the Anr regulon (113), the Ppk2 and the AdhA alcohol dehydrogenase while NorC was not detected in HACs but present in NACs and the ancestral strain. Only three proteins from the Anr core regulon being altered is surprising, since hypoxia response has been primarily associated with it. However, as previously mentioned, Anr regulation is complex, and hypoxia adaptation may have selected for strains that downregulate Anr to conserve energy. Additionally, as mentioned the moderate hypoxic conditions utilised may lead to adaptation via other pathways rather than the Anr regulon. Notably, the Anr regulon shares homologous genes with the DosR regulon of *Mycobacterium abscessus* and Lxa locus of *Burkholderia cenocepacia*, highlighting that aspects of the hypoxia response may be shared across several opportunistic CF lung infections more generally. Investigations on the impact of different levels of hypoxia on these three pathogens is ongoing in our group. Targeting shared mechanisms of adaptation to chronic infection could present a new opportunity for the treatment of multiple pathogens simultaneously.

### Impact

Overall, we demonstrate that alterations in phenotype after long-term exposure to hypoxic conditions shows several similarities associated with chronic infection. Further understanding of the mechanisms by which *P. aeruginosa* adapts to hypoxia in chronic infection and the interplay with other factors present in the lung environment will potentially enable us to prevent or treat the switch from acute to chronic infection. This could inevitably limit antibiotic use, severity of the disease and ultimately, decrease the socioeconomic impact of infections caused by *P. aeruginosa*. Furthermore, the first drug to alter the human response to hypoxia was recently approved for use by the US Food and Drug Administration (175). Thus, it is tempting to speculate that the response of bacteria to low oxygen pressures could be targeted to prevent a switch from acute to chronic infection.

The observed changes in proteome and phenotypes (increased biofilm formation, increased antibiotic resistance, decreased siderophore production, increased SCV prevalence) further deepens our knowledge on how hypoxia leads to adaptation of *P. aeruginosa*, shedding light on how prolonged low oxygen exposure can impact on adaptation to chronic infection. Overall, we confirm that hypoxia contributes to the development of multiple phenotypic changes that are prevalent in chronic infection, opening a path towards designing therapies targeting these.

## Materials and methods

### Bacterial strains

*Pseudomonas aeruginosa* AMT 0023-30 early infection clinical strain (E) and AMT 0023-34 late infection clinical strain (L) are two well characterised strains in the International *P. aeruginosa* panel (35). Hypoxia- and normoxia-adapted *P. aeruginosa* strains were obtained in this study as described below and are referred to as H1, H2, H3, N1, N2, and N3 (Table 1). If not stated otherwise, bacteria were routinely grown in LB broth (Sigma-Aldrich, St. Louis, MO, USA) at 37°C with orbital agitation (200 rpm) or plated on LB agar (Sigma-Aldrich, St. Louis, MO, USA) and grown at 37°C.

### Experimental evolution

Overnight cultures of *P. aeruginosa* strain AMT 0023-30 (E) were inoculated into 30 ml LB media to OD_600_ ∼0.1. Prior to use, the media was equilibrated for 18 h at 37°C in an O_2_ Control InVitro Glove Box (PHGB model, Coy Laboratory Products, MI, United States) at 6% O_2_, 5% CO_2_ or in a Steri-Cycle CO_2_ Incubator (model 371, Thermo Scientific, Waltham, MA, USA) at 21% O_2_, 5% CO_2_. Three flasks were inoculated as biological replicates for each of the conditions being examined. Additionally, aliquots (2 ml) of the overnight *P. aeruginosa* cultures were retained as a control for proteomic and transcriptomic analysis. All flasks were incubated for 24h with agitation (200 rpm, CO_2_ resistant orbital shaker, model 15351105, Thermo Fisher Scientific, Waltham, MA, USA) in hypoxic and normoxic conditions. Daily aliquots (2 ml) were removed for proteomic analysis from each of the flasks and also additional aliquots retained for OD_600_ measurements and plating. Glycerol stocks (50%, LB (v/v)) were prepared for subsequent phenotype analysis and frozen immediately to ensure minimal oxygen exposure. A volume of 5% of the bacterial culture was also transferred into 30 ml of pre-equilibrated media each day and this procedure was continued for up to 28 days to determine long-term adaptation patterns. To ensure stability of the hypoxic conditions, all of the samples from H1, H2 and H3 were taken inside the hypoxia chamber. To ensure that there were no other environmental differences between the test conditions other than oxygen levels, the same batch of media was used for both normoxia-adapting and hypoxia-adapting flasks and shaking conditions were consistently maintained.

### Whole proteome analysis

Samples (2 ml) were collected throughout the experimental evolution for proteome analysis. Bacterial cells were pelleted at 14000-16000 g for 2 min and stored at -80°C until further analysis. Whole cell lysates were extracted from each of the hypoxia- and normoxia exposed cultures by resuspending in 1 ml of ice-cold lysis buffer containing 40mM Tris-HCl (pH 7.8) and 1x cOmplete™, EDTA-free protease inhibitor cocktail (Roche, Basel, Switzerland). Cells were disrupted by a sonication with a sonic dismembrator and microtip 6mm probe (model 505; Fisherbrand, Thermo Fisher Scientific, Waltham, MA, USA) (set with 20% output power using eight 10 s bursts on ice). Cell debris and any remaining intact cells were pelleted using centrifugation (12000 g for 30 min). Supernatants were transferred into fresh tubes and the proteins were precipitated with four volumes of methanol and incubated in -80°C overnight. The samples were centrifuged at 12000 g for 30 min at 4°C, supernatants were removed, and the protein pellets were resuspended in 200 μl of resuspension buffer containing 8 M urea, 50 mM Tris-HCL (pH 8.0), before 1 M 1,4-dithiothreitol (DTT) was added to each supernatant (10 μl/ml lysate) and incubated at 56°C for 30 min, followed by the addition of 1 M iodoacetamide (55 μl/ml lysate) which was incubated in darkness at room temperature for 20 min. Lysates were dialyzed in SnakeSkin™ tubing (Thermo Fisher Scientific, Waltham, MA, USA) with a cut-off of 3.5 kDa against 100 mM ammonium bicarbonate overnight, with stirring at 4°C, followed by at least 4 more hours incubation with fresh volumes of ammonium bicarbonate. Trypsin (400 ng/μl) (Sigma-Aldrich, St. Louis, MO, USA) was added to the dialyzed protein (5 μl /100 μl protein) and tubes were incubated overnight at 37°C. Aliquots from each sample were placed in fresh tubes and samples were dried in a vacufuge concentrator system (model 5301; Eppendorf, Hamburg, Germany) at 65°C and resuspended in 20 μl of ZipTips resuspension buffer containing 0.5% trifluoroacetic acid (TFA) in MilliQ water. ZipTips (Merc Millipore, Burlington, MA, USA) were used for peptide purification as per the manufacturers’ instructions. Nanodrop spectrophotometer DS-11+ (DeNovix, Wilmington, DE, USA) was used for peptide concentration measurements. The samples were analysed on a Bruker TimsTOF Pro mass spectrometer connected to a Evosep One chromatography system. Peptides were separated on an 811cm analytical C18 column (Evosep, 3 µm beads, 100 µm ID) using the pre-set 33 samples per day gradient on the Evosep one. A trapped ion mobility (TIMS) analyser was synchronized with a quadrupole mass filter to enable the highly efficient PASEF (Parallel Accumulation Serial Fragmentation acquisition) procedure with acquisition rates of 100 Hz. The accumulation and ramp times for the TIMS were both set to 100 ms., with an ion mobility (1/k0) range from 0.62 to 1.46 Vs/cm. Spectra were recorded in the mass range from 100 to 1,700 m/z. The precursor (MS) Intensity Threshold was set to 2,500 and the precursor Target Intensity set to 20,000. Each PASEF cycle consisted of one MS ramp for precursor detection followed by 10 PASEF MS/MS ramps, with a total cycle time of 1.16 s.

Protein identification and label-free quantitative (LFQ) analysis were conducted using MaxQuant (v 2.4.2.0, https://maxquant.org/) by searching against the reference strain *P. aeruginosa* PAO1 (ATCC 15692). For protein identification, the following search parameters were used: trypsin was the digesting enzyme with up to two missed cleavages allowed for; oxidation on methionine and acetylation on the N terminus were selected as variable modifications with a fixed modification of carbamidylation on cysteine; Label Free Quantitation (LFQ) and Match Between Runs (MBR) options were selected. FDR for protein and peptide were set at 1%.

The three biological replicates from each 28 day culture condition were analysed, with one technical replicate for a randomly chosen sample and LFQ intensities were compared. All missing LFQ intensities were imputed with a fixed value of 1. The mean was calculated separately for the hypoxia and normoxia samples, followed by fold change (FC) calculation.

One-way ANOVA statistical analysis (p-value ≤0.05) together with Benjamini–Hochberg false-discovery adjustment were performed to identify proteins with a significantly changed abundance Proteins with adjusted p-values≤0.1 were considered significantly altered in abundance.

Functional enrichment of proteins significantly changed in abundance was performed. For GO biological process functional enrichment analysis ShinyGO (v 0.82) (44) was used. Species was set to *P. aeruginosa* PAO1, FDR cutoff was set to 0.05 and the minimal pathway size was set as two. For STRING local network and KEGG pathway enrichment the STRING (v 12.0) database was used with default settings.

### Antibiotic resistance

Overnight bacterial cultures were diluted in 10 ml of fresh LB to OD_600_ equal to 0.1. Bacteria were spread evenly on Mueller–Hinton agar plates (Neogen, Lansing, MI, USA) using cotton swabs, in three directions, to give a homogenous lawn. Antibiotic disks (Oxoid, Hampshire, UK) were placed on the surface of the agar and the plates were incubated overnight in normoxic conditions at 37°C. Growth inhibition zone diameters were measured in at least two points, and the mean was used for statistical analysis. The experiment was repeated three times. The following antibiotics were tested: norfloxacin (10 µg), ciprofloxacin (10 µg), azithromycin (15 µg), clarithromycin (15 µg), ceftazidime (30 µg), meropenem (10 µg), imipenem (10 µg), piperacillin (100 µg), aztreonam (30 µg), tobramycin (10 µg), and tetracycline (30 µg).

### Motility

Motility assays were performed as described previously (176). Swimming motility was assessed by inoculating the surface of a 0.3 % (w/v) LB agar plate with overnight culture using a sterile toothpick. Swimming and swarming plates were incubated overnight in normoxic conditions at 37°C for ∼18 h. For swarming motility 0.8% nutrient broth (Sigma-Aldrich, St. Louis, MO, USA), 0.5% dextrose, 0.5% agar plates were inoculated identically as for swimming motility assessment. The diameter of bacterial growth zones was measured in at least two directions, and the mean was used for statistical analysis. Twitching motility was assessed by inoculating 1.5 % (w/v) LB agar plate by piercing through the agar. Plates were incubated overnight in normoxic conditions at 37°C for ∼18 h, and the agar was removed. Bacteria on the surface of the plate were stained with 1% (w/v) crystal violet for 20 min. The plates were washed with water until all excess dye was removed and dried at room temperature. The diameter of the growth zone was measured in at least two points, and the mean was used for statistical analysis. The experiment was repeated three times.

### Biofilm production

Overnight cultures of *P. aeruginosa* strains were diluted in fresh LB media to OD_600_ of ∼0.1, and 100 µl of each diluted culture was transferred into 6 wells of a 96-well plate. The plates were incubated statically in normoxic conditions at 37°C for 24, 48 or 72 h and OD_600_ was measured with the Synergy H1 microplate reader (Mason Technology, Dublin, Ireland). The medium with planktonic bacteria was removed, and the biofilm fixed by incubation with 200ul of methanol for 10 min. The wells were washed three times with phosphate buffered saline (PBS) (Sigma-Aldrich, St. Louis, MO, USA), followed by the addition of 200 µl of 0.1% crystal violet solution and 30 min incubation at room temperature. The solution was removed, and the wells were washed three times with water and once with PBS. The plates were dried for at least 30 min at room temperature and 150 µl of 96% ethanol was added to dissolve the bound stain. After 10 min of incubation at room temperature, the solution was mixed by pipetting, OD_590_ was determined, and the OD_590_/OD_600_ ratio was calculated. The experiment was repeated three times.

### Exopolysaccharide production - Congo Red agar assay

Overnight cultures of *P. aeruginosa* strains were pelleted and washed with 1 ml of PBS. The pellets were resuspended in PBS to OD_600_ of ∼0.1 and 1 µl of the suspension was plated on a Congo Red agar plate (10 g/L Tryptone, 10 g/ L Bactoagar, 40mg/L Congo Red, 20mg/L Coomassie Brilliant Blue). The plates were incubated for 48 h in normoxic conditions and colony morphology was assessed. The experiment was repeated three times.

### Stress sensitivity assays

Overnight cultures of *P. aeruginosa* strains were diluted to a CFU/ml of 10^8^ and a range of serial dilutions was prepared. Suspensions (1µl) were plated on one of the following: LB agar control plates, LB agar supplemented with 0.5 M NaCl (osmotic stress), LB agar supplemented with 1mM H_2_O_2_ (oxidative stress), or LB agar, pH 5 (acidic stress). The plates were incubated in normoxic conditions at 37°C for 24 or 48 h. Temperature stress was determined by incubation of an LB agar plate at 45°C for 24 h. Change of sensitivity was determined by determining the lowest dilution at which bacterial growth was observed. The experiment was repeated three independent times.

### Galleria mellonella infection model

Virulence was determined as previously described (37). Wax moth larvae (*Galleria mellonella*) (The KammieShop, Waalwijk, Denmark) were equilibrated in normoxic conditions at 15°C for at least 7 days after arrival and stored at the same conditions for up to 4 weeks, during which larvae with a weight range of 0.2-0.4 g were selected for experiments. Overnight bacterial cultures in LB were diluted in fresh media to OD_600_ of ∼0.111and grown for 2-3 h. Cultures were centrifuged, and bacterial cells were washed with PBS. The bacteria were diluted to 10^6^ CFU/ml and serially diluted to 10^−116^. Each dilution was injected (20 μl) into the hindmost left proleg of healthy larvae (10 per group) and the same volume of PBS was injected into one group as a control. Injected larvae were incubated at 37°C for 72 h. To have an accurate count of the bioburden inoculated, aliquots of the bacteria injected were serially diluted and plated onto LB agar and colonies counted after 2411h. The experiment was repeated at least three times for each strain tested.

### Pyocyanin production quantification

*P. aeruginosa* strains were inoculated to OD_600_ in fresh PB media (20g/L Bacto-Peptone, 1.4g/L MgCl_2_, 10g/L K_2_SO_4_) and cultured with aeration in normoxic conditions at 37°C to maximize pyocyanin production. The bacterial cultures were pelleted (10 min at 12300 g) and two 7.5-ml replicates were withdrawn from each of the 18 h stationary phase cultures. Pyocyanin was extracted with 4.5 ml of chloroform and a total of 1.5 ml of 0.2 M HCl was added to each extract causing colour change from blue to pink. The absorbance of this solution was measured at 520 nm, and the obtained values were converted to pyocyanin content following Essar et al. (1990) (48). The experiment was repeated at least two times.

### Siderophore production

The siderophore production was quantified with a modified version of a microplate method developed previously (177). Prior to siderophore quantification all glassware was rinsed with a 3M solution of HCl to remove trace iron and rinsed thoroughly with deionised water. The CAS reagent was prepared as described before (177).

Three independent cultures of each tested population or strain were grown in fresh LB medium for 48 h at 37°C with agitation in normoxic conditions. Aliquots of the cultures were taken for CFU counts, serially diluted and plated on LB agar. The cultures were spun down (10 min at 12300 g), and 50 µl aliquots of the supernatant were transferred into separate wells of a microplate followed by an immediate addition of 100 µl of CAS reagent. As a control LB broth was used. The plates were incubated for 20 min at room temperature in the dark and the optical density was measured at 630 nm using the Synergy H1 microplate reader. The Siderophore units were calculated as follows: siderophore production = [(A_LB_ - A_s_) * 100]/A_LB,_ where A_LB_ = absorbance of LB control and A_s_ = absorbance of sample. The earlier plated bacteria were incubated at 37°C overnight and colony forming units were counted. Siderophore production per colony forming unit was calculated to account for any differences in growth. The experiment was repeated two times.

### Determination of growth by optical density

To assess the influence of the inoculation volume on the growth patterns of *P. aeruginosa* AMT 0023-30, 24h stationary phase cultures were inoculated into fresh LB media in a way they constituted 1%, 5% or 10% of the new culture. Three 200 µl technical replicates of the cultures were transferred into a 96-well plate and incubated in the Synergy H1 microplate reader in normoxic conditions at 37°C with agitation with OD_600_ measurements taken every 30 min.

### Determination of growth by CFU

To assess the differences in the numbers of generations in hypoxic conditions and normoxic conditions of the AMT0063-30 early infection strain overnight cultures were prepared. The following day they were inoculated in triplicate into fresh LB media and incubated with aeration at 37°C in the O_2_ Control InVitro Glove Box in 6% O_2_, 5% CO_2_ or in 21% O_2_, 5% CO_2_ or in the Steri-Cycle CO_2_ Incubator at 21% O_2_, 5% CO_2_. Every hour a sample was taken from each culture for OD_600_ measurements and plating on LB agar. The agar plates were incubated overnight at 37°C and the colony forming units were calculated and OD_600_ to CFU/ml was plotted, a trend line was determined and R^2^ calculated. The doubling time was calculated for the logarithmic phase of growth using the following formula: Doubling time = (Duration (min)\**ln*(2))/[*ln*(final cell concentration/initial cell concentration)].

### Intracellular c-di-GMP concentration measurements

C-di-GMP was extracted from *P. aeruginosa* cells as previously described (178). Briefly, the OD₆₀₀ of overnight cultures was measured, and a culture volumes equivalent to 4 ml of OD₆₀₀ = 1.8 were centrifuged and washed twice with PBS. Pellets were resuspended in 400 µl PBS, incubated at 100°C for 5 min before adding ice-cold ethanol (final concentration of 65%), and mixing. The supernatants were collected after centrifugation and the extraction was repeated three times. Pooled supernatants were dried at 30°C, resuspended in 200 µl nanopure water, and c-di-GMP levels were quantified using a cyclic-di-GMP assay kit (Lucerna Technologies, Brooklyn, NY, USA), as per the manufacturer’s recommendations. C-di-GMP concentrations were normalized to total protein content, measured using the Bradford reagent (Sigma-Aldrich, St. Louis, MO, USA) on pellets that were resuspended in TE buffer, sonicated on ice for 1 min (10 s bursts, 20% output power), centrifuged and the supernatants used for protein quantification.

### Statistical analysis of phenotype data

Initial data preparation was performed using Microsoft Excel and statistical analysis was performed using GraphPad Prism (v 8.0.2, GraphPad Software, Boston, MA, US). Only p-values of 0.05 or lower were considered statistically significant. First, the normality was assessed using the Shapiro-Wilk test. Data with normal distribution was analysed using the Ordinary one-way ANOVA with the Uncorrected Fisher’s LSD; data with non-normal distribution was analysed using the Kruskal-Wallis test with an uncorrected Dunn’s test. Details of the analysis and significance are provided in figure legends.

## Acknowledgements

JD, CJC are supported by an SFI Frontiers of the Future award (20/FFP-P/8717) to SMcC. ND was supported by the Irish Research Council, GOIPG/2021/1443 IRC Award. The authors are grateful to Dr David Gomez-Matallanas for his advice in proteomic analysis and for the support of the staff and facilities in the UCD Conway proteomics core.

## Supporting information captions

S1. Figure. Differences in growth patterns of the P. aeruginosa AMT 0023-30 early infection strain with different inoculation volumes.

S2. Figure. Differences in doubling times of the P. aeruginosa AMT 002330 early infection strain in hypoxic and normoxic conditions. A. OD_600_ measurements over time. B. CFU/ml counts over time C. OD_600_ to CFU/ml trend lines and linear regression calculations. D. Differences in generation times in normoxia and hypoxia.

S3. Figure. Stability of SCVs in normoxic conditions. A. Photograph of different P. aeruginosa colony variants streaked on LB agar and grown overnight in normoxic conditions. B. Enlarged fragment of photograph in section A.

S4. Table. Proteomic analysis of long-term hypoxia-adapted P. aeruginosa isolates. A. All proteins differentially expressed between HACs and the ancestral strain (D0), NACs and the ancestral strain (D0) or HACs and NACs. B. Proteins not detected in at least two replicates in at least one of the groups (Day 0, HACs, NACs). C. All proteins detected in the proteomics analysis of the ancestral strain, HACs and NACs

S5. Table. Functional enrichment of the 89 proteins of interest significantly altered in abundance after long-term adaptation of P. aeruginosa to hypoxia

S6. Figure. Effects of long-term hypoxia adaptation on intracellular c-di-GMP concentrations. The graph represents the concentration of c-di-GMP in the cell extract from an equivalent of 4 ml of culture of OD_600_ =1.8. No statistically significant differences between cultures were shown with the Brown-Forsythe ANOVA test. E: early infection clinical strain (AMT 0023-30); L: late infection clinical strain (AMT 0023-34); N1, N2, N3: 28-day normoxia-adapted populations; H1, H2, H3: 28-day hypoxia adapted populations.

## Notes

### Competing Interest Statement

The authors have declared no competing interest.

### Summary of Updates

This revision provides a reanalysis of the proteome data to compare hypoxia adapted cultures directly with the ancestral strain, in addition to the comparison with normoxia adapted cultures.

