## Supplementary figures and images for "Long-term adaptation to hypoxia provides insight into mechanisms facilitating the switch of *Pseudomonas aeruginosa* to chronic lung infections"

### Supplemental figures S1, S2, S3 and S6

#### Slide 1
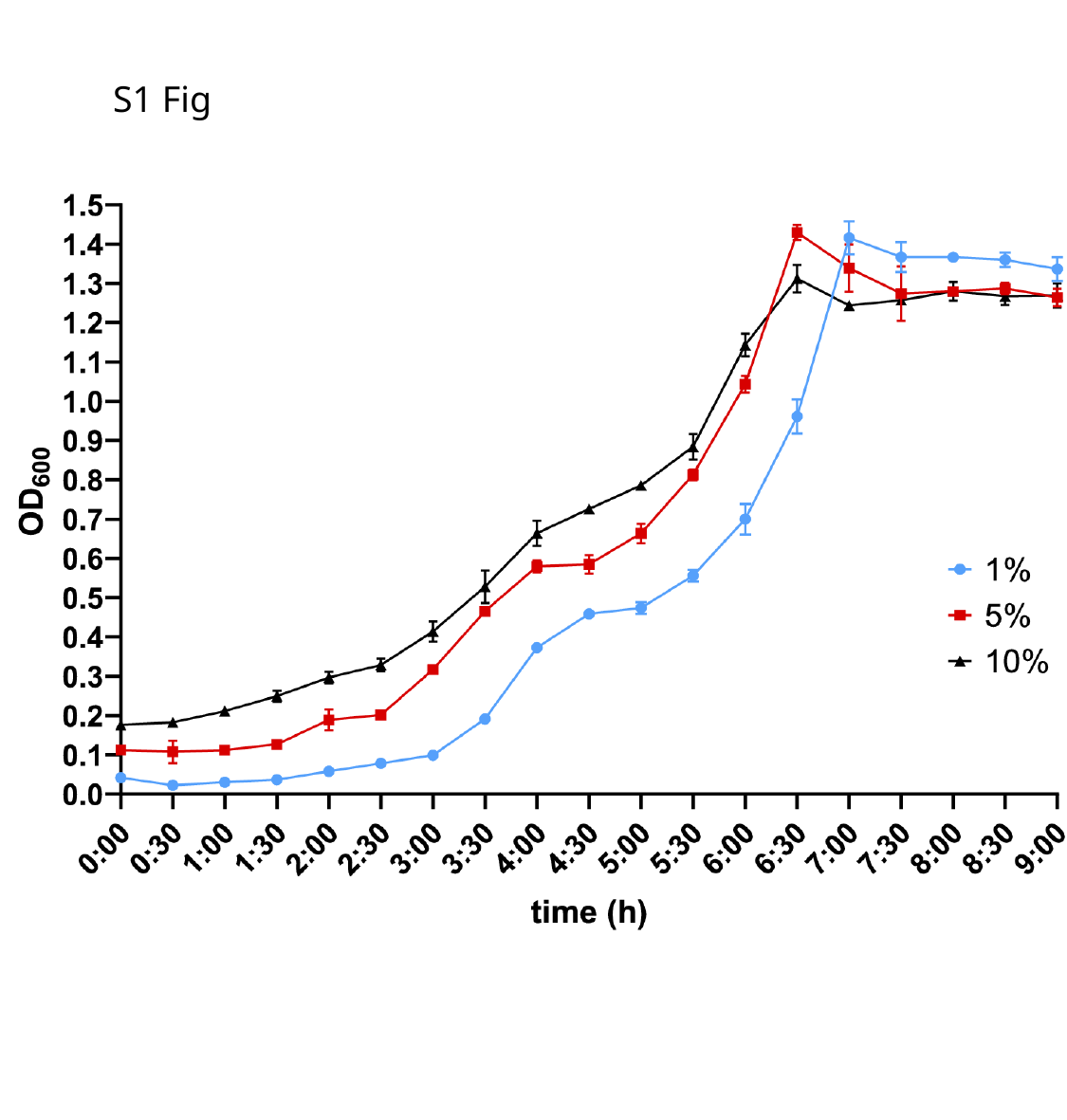

S1 Fig

#### Slide 2
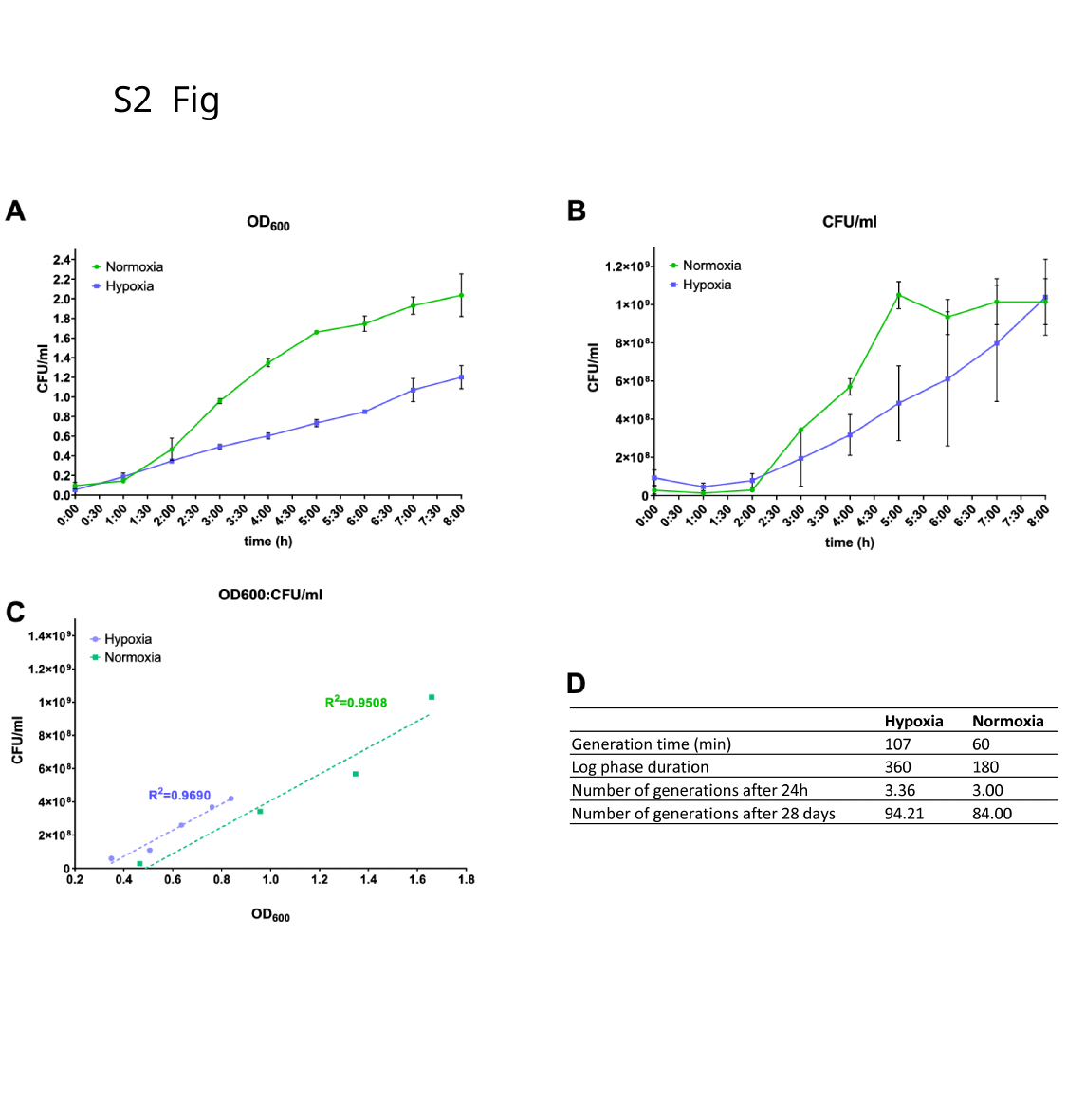

S2 Fig

#### Slide 3
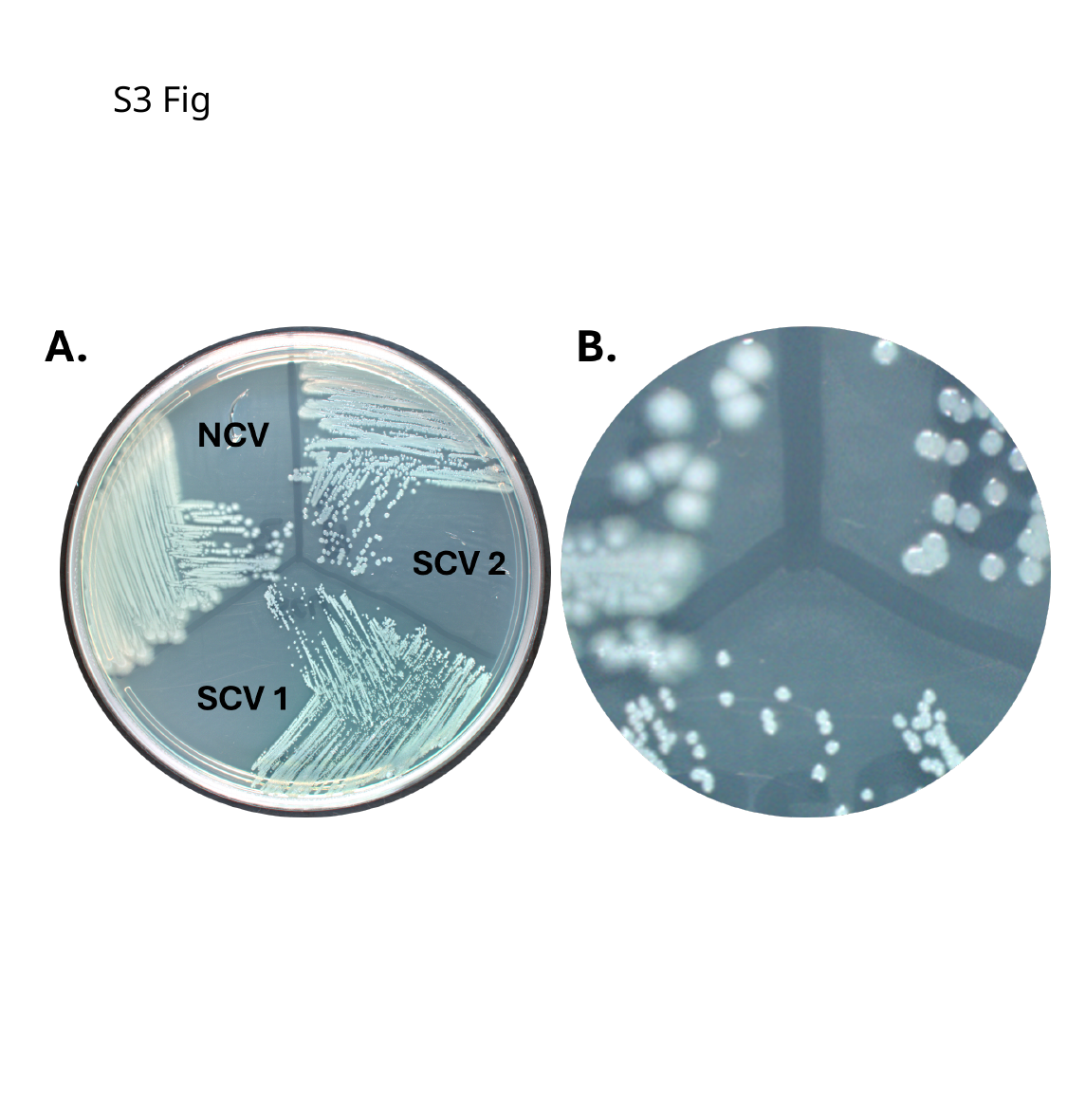

S3 Fig

#### Slide 4
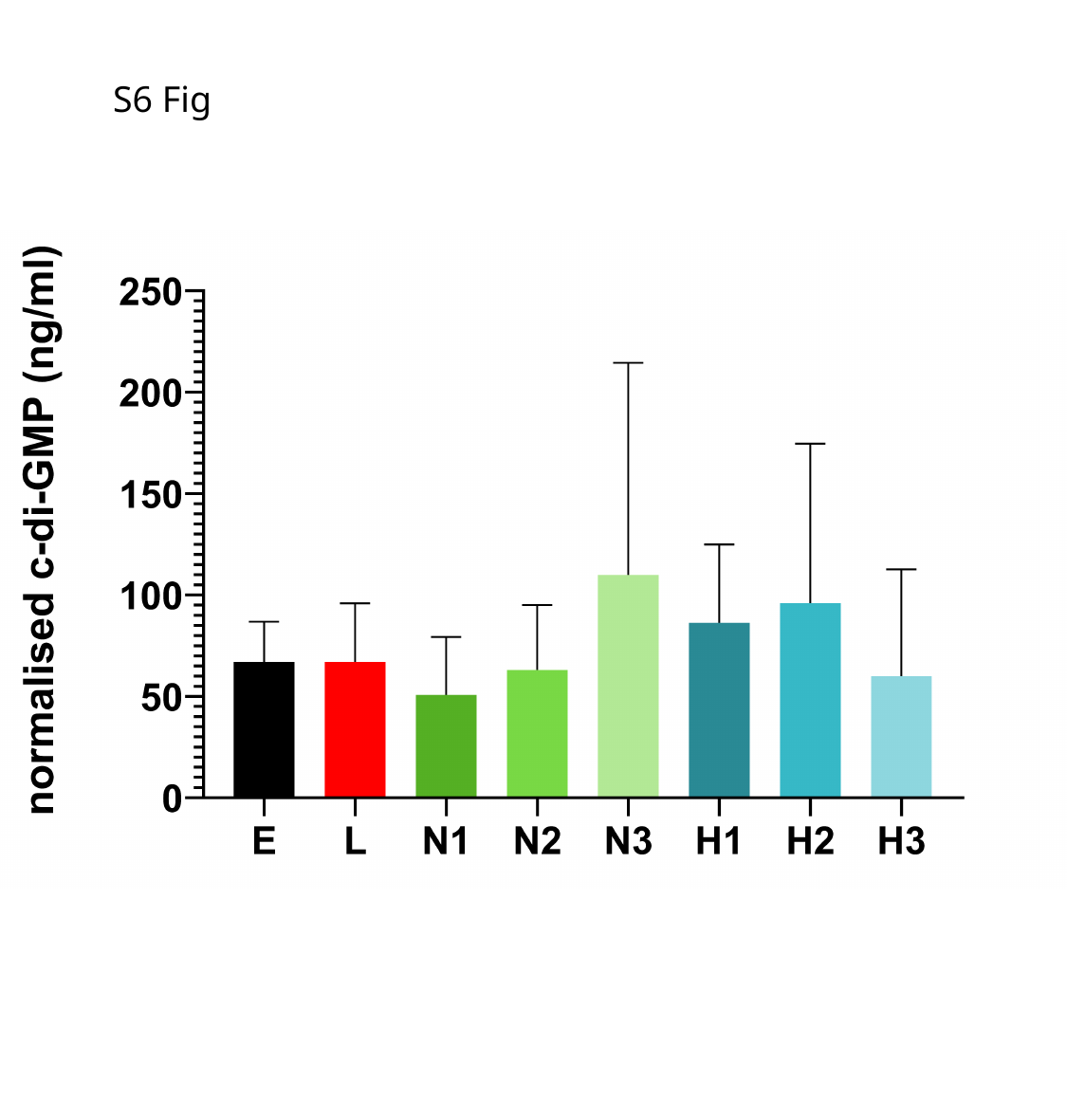

S6 Fig
